# “Personalized Brain Morphometric Feature as a Transdiagnostic Predictor of Psychopathology: Insights from Dual Systems Models”

**DOI:** 10.1101/2025.04.14.644899

**Authors:** Min-fang Kang, Kun-ru Song, Jia-lin Zhang, Lin-xuan Xu, Ya-jie Zhang, Zi-heng Zhao, Hui-ying Deng, Xiao-yi Fang, Marc N. Potenza, Jin-tao Zhang

## Abstract

**Background:** Adolescents are particularly vulnerable to developing psychopathological symptoms, yet neurobiological markers that can identify these vulnerabilities in a personalized and interpretable manner remain limited. Dual Systems Models suggest that this vulnerability may result from asynchronous development of neural systems subserving cognitive-control and socioemotional functions. Given that rigorous empirical evidence is sparse, this study aimed to quantify developmental imbalances and evaluate their predictive value for psychopathology.

**Methods:** Based on Dual Systems Models, the dorsolateral prefrontal cortex (DLPFC) and ventral striatum (VS) were selected as key regions for the two brain systems, respectively. Using longitudinal data from the Adolescent Brain Cognitive Development (ABCD) study (baseline: n=11,238, ages 9.92±0.625 years; 2-year follow-up: n=7,870, ages 12.2±0.652 years; 4-year follow-up: n=2972, ages 14.1±0.693 years), we derived a personalized imbalance score based on brain morphometric features, including surface area, thickness and gray-matter volume. Nested linear mixed models were employed to assess associations between the imbalance scores and psychopathology. Additionally, generalized additive models were used to capture developmental trajectories of the imbalance scores and explore potential non-linear associations with psychopathology. To examine reproducibility, data from the Lifespan Human Connectome Project in Development study (HCP-D, N=652, ages 8-21 years) were interrogated.

**Results:** The imbalance score, quantified as the difference between VS volume and DLPFC surface area, exhibited strong reliability and validity. During adolescence, the score exhibited a declining trend, with decreasing growth rates and variability between individuals. The score was positively associated with externalizing symptoms and showed a U-shaped relationship with internalizing symptoms. These findings were replicated in the HCP-D sample.

**Conclusions:** The novel neuroanatomical imbalance score empirically supports Dual Systems Models and may function as a transdiagnostic marker for internalizing and externalizing psychopathology. These findings enhance understanding of the neurodevelopmental mechanisms underlying adolescent psychopathology and offer potential implications for precision prevention and intervention strategies.

## Introduction

Adolescence is a critical developmental period characterized by profound physical, psychological, and neurobiological changes, alongside heightened vulnerability to psychiatric disorders (1,2).

These disorders, which manifest through diverse psychopathological symptoms, are both highly prevalent and related to significant societal and personal consequences, necessitating the development of effective preventive interventions for youth (3). However, progress has been hindered by etiological heterogeneity and a lack of precise, interpretable measures for quantifying neurobiological dysfunctions. While data-driven “black box” approaches have gained traction, their limited interpretability restricts their utility. In contrast, theory-driven approaches provide a more neurobiologically grounded framework for identifying mechanistic substrates (4). Within a neurodevelopmental framework, personalized neuroimaging-derived measures offer the potential to enhance early detection of psychopathological symptoms, enabling the identification of high-risk subpopulations, which could facilitate targeted interventions and ultimately reduce healthcare burdens (5).

Psychopathological symptoms peak during adolescence (6), as highlighted by multiple neurodevelopmental theories that aim to explain this vulnerability (7,8), including the Dual Systems Model (9), Maturational Imbalance Model (10), and Driven Dual Systems Model (11). These frameworks converge on a central premise: heightened vulnerability during adolescence stems from the asynchronous development of two key brain systems – a socioemotional system, centered in subcortical regions especially the ventral striatum (VS), and a cognitive-control system, centered in the dorsolateral prefrontal cortex (DLPFC) (12). The socioemotional system matures earlier, increasing sensitivity to internal emotional states and enhancing the propensity for external risky behaviors—both of which are associated with psychopathological symptoms. In contrast, the slower-maturing cognitive-control system is not yet fully developed during adolescence, limiting its capacity to effectively regulate internal and external factors (13). Although studies may use different terminologies, we refer to this asynchronous development as “*dual-systems imbalance”* (14,15), which underscores the dynamic interplay of individual differences and reflects the relative dominance of one system over the other. For consistency, we refer to these theories as “*Dual Systems Models.*”

While many studies have focused on population-level patterns, Dual Systems Models emphasize the importance of individual differences, highlighting that not all adolescents develop psychopathological symptoms. This variability necessitates exploration of why some individuals are more vulnerable than others (10,13). Neurodevelopmental processes, marked by significant changes in brain morphology, related to synaptic pruning, myelination, and cortical expansion, refine motor, cognitive, and regulatory functions during adolescence (16), which may drive individual differences in dual-systems imbalance.

Dual-systems imbalance has been linked to several psychiatric conditions including adolescent depression and anxiety (7,8,17,18), categorized within the internalizing dimension of the Hierarchical Taxonomy of Psychopathology (HiTOP) framework (https://www.hitop-system.org/) (19,20). The HiTOP framework integrates dimensional structures of maladaptive personality and psychiatric disorders into a hierarchical structure encompassing broad spectra, including internalizing and externalizing dimensions (21). Additionally, dual-systems imbalance has been linked to aggression, conduct disorder, and substance use (22–27), which fall within the HiTOP externalizing dimension. These findings suggest that dual-systems imbalance may serve as a transdiagnostic predictor, capturing shared variance across internalizing and externalizing dimensions of psychopathology.

Despite considerable empirical support for Dual Systems Models (13), several gaps remain. Anatomical measures have proven valuable in revealing individual differences, yet neuroimaging studies often focus on single brain morphological features rather than integrating multiple metrics, such as surface area (SA), surface thickness (ST), cortical volume (CV) and subcortical volume (SV), which independently capture informative neuroanatomical differences (28,29). To maximize the predictive value of brain morphometry and enhance the validity and clinical relevance of dual-systems imbalance, a comparative analysis to identify optimal neural correlates is indicated.

Second, specific conceptualizations of dual-systems imbalance have received limited attention, hindering direct and comprehensive testing. Much of the existing research on Dual Systems Models has focused on analyzing the two systems separately, often using categorical comparisons to examine their individual trajectories. For instance, prior studies have explored individual effects of heightened sensation-seeking and low self-regulation, two psychological constructs central to dual-systems, on adolescent risk-taking (Fig 1B). However, how these systems interact with each other has often been overlooked (8,9,12). This approach fails to directly capture the concept of “imbalance” as examining independent effects and does not consider the relative standing of one system in relation to the other, thereby providing no insight into the imbalance itself. To address this limitation, we propose integrating the two systems into a single, comprehensive index that offers a direct and clear representation of imbalance, creating new opportunities to operationalize and quantify dual-systems imbalances across adolescence.

**FIGURE 1.**
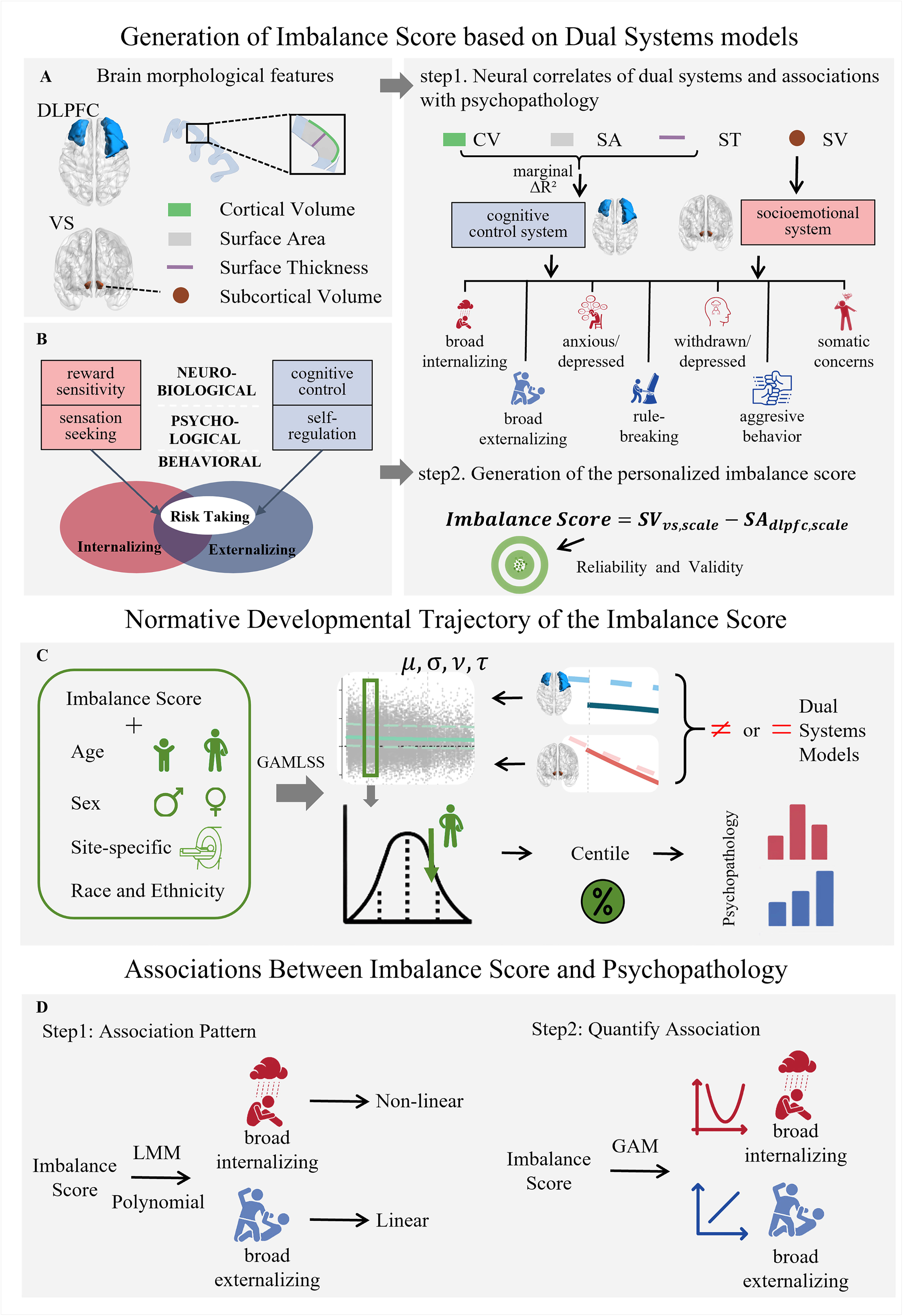
Outline of the study ***Panel A.*** Brain phenotypes were measured in surface-based morphometry. We defined the regions of interest from the meta-analytic platform Neurosynth and refined them based on quality control. ***Panel B.*** The constructs implicated in the Dual Systems Models (13). ***Panel C.*** GAMLSS modeling was used to estimate the relationship between personalized imbalance score and age, and controlling for sex, race, ethnicity and variation between scanning sites. The normative trajectory of the median for imbalance score was plotted as a population reference curve. Abbreviations: DLPFC, dorsolateral prefrontal cortex; VS, ventral striatum; SA, surface area; SV, subcortical volume; ST, surface thickness; CV, cortical volume; LMM, linear mixed model; GAM, generalized additive model; GAMLSS, generalized additive models for location, scale and shape

To understand adolescent psychopathology, it is important to assess the developmental trajectory of this imbalance. Understanding not only whether the imbalance changes with age but also the nature and trajectory (including shape and form) of these changes is fundamental to developmental science (30). Defining the shape and form of the imbalance has important implications for understanding mechanisms of neuroplasticity and identifying factors influencing adolescent critical periods (31). However, prior studies have relied on fixed developmental shapes (e.g., linear or quadratic trajectories) or categorical comparisons (e.g., adolescents vs. adults) (12,32), limiting the ability to examine the dynamic interplay between these systems, the rate of change, and the timing of stability—key elements for precise developmental science (31). Such approaches have also hindered the resolution of theories, where distinct linear and non-linear shapes have been proposed. Advances in statistical methods, such as generalized additive models for location, scale, and shape (GAMLSS), now allow for more precise modeling of potentially non-linear developmental trajectories, addressing prior challenges in understanding normative developmental processes.

Taken together, we introduce a novel, individual-level integrated measure (hereafter refer to as the “*imbalance score*”) to quantify the dual-systems imbalance during adolescent development, using multiple brain morphometric features. Morphometric measures were selected for their stability, state-independence, and capacity to capture individual differences (Supplement 7). Based on the HiTOP framework, we investigated associations between the imbalance scores and two key psychopathological dimensions: internalizing and externalizing problems. These analyses utilized a large longitudinal sample from the Adolescent Brain and Cognitive Development (ABCD) Study and validated from an independent sample from The Lifespan Human Connectome Project in Development (HCP-D). We hypothesized that: (1) the individual-level imbalance score would demonstrate high reliability and validity; (2) the normative trajectory of imbalance scores would align with the heuristic theory, with atypical scores predicting greater psychopathology; and (3) the imbalance scores would correlate with both internalizing and externalizing problems, supporting its potential transdiagnostic potential.

## Methods and Materials

### Participants and Data Preprocessing

The ABCD study is an ongoing longitudinal project that recruited at its onset 11,868 children aged 9-10 years from 21 U.S. sites (33). This study aims to characterize psychological and neurobiological development from preadolescence to emerging/young adulthood. Behavioral and neuroimaging data from the ABCD 5.1 data release were analyzed, involving baseline (T0, n = 11,238, 5412 girls, mean age = 9.92±0.625 years), 2-year follow-up (T2, n = 7,870, 3662 girls, mean age = 12.2±0.652 years), and 4-year follow-up (T4, n = 2972, 1399 girls, mean age = 14.1± 0.693 years) waves. Sample selection is summarized in Table S1, and the demographic characteristics are provided in Table 1. To assess generalizability, data from the HCP-D study were analyzed. The cross-sectional HCP-D, involving participants aged 5 to 21 years with embedded longitudinal cohorts around puberty, collected data across four US sites (34). Neuroimaging and psychopathology data from the HCP-D 2.0 release were available for 591 participants aged 8–21 years.

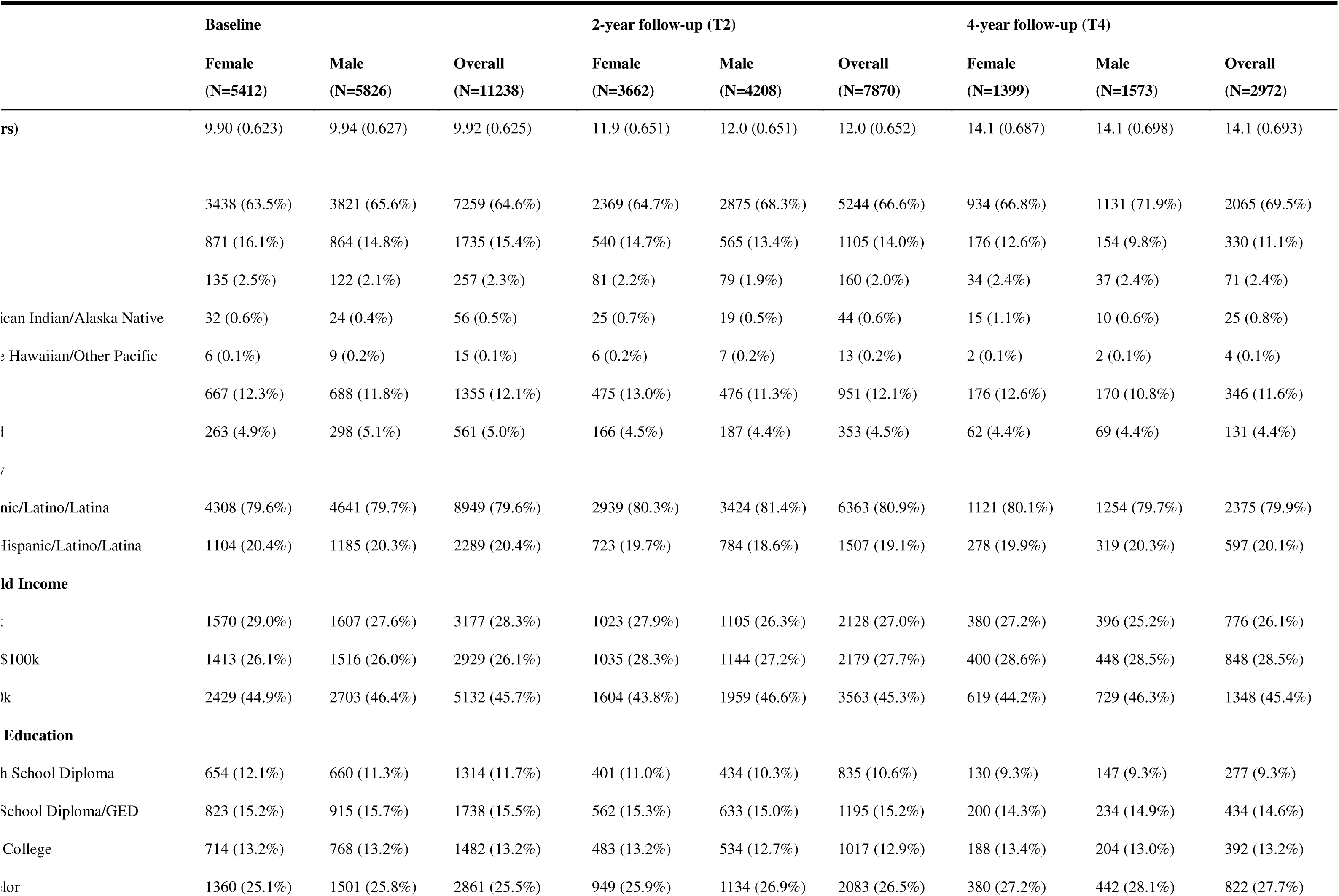
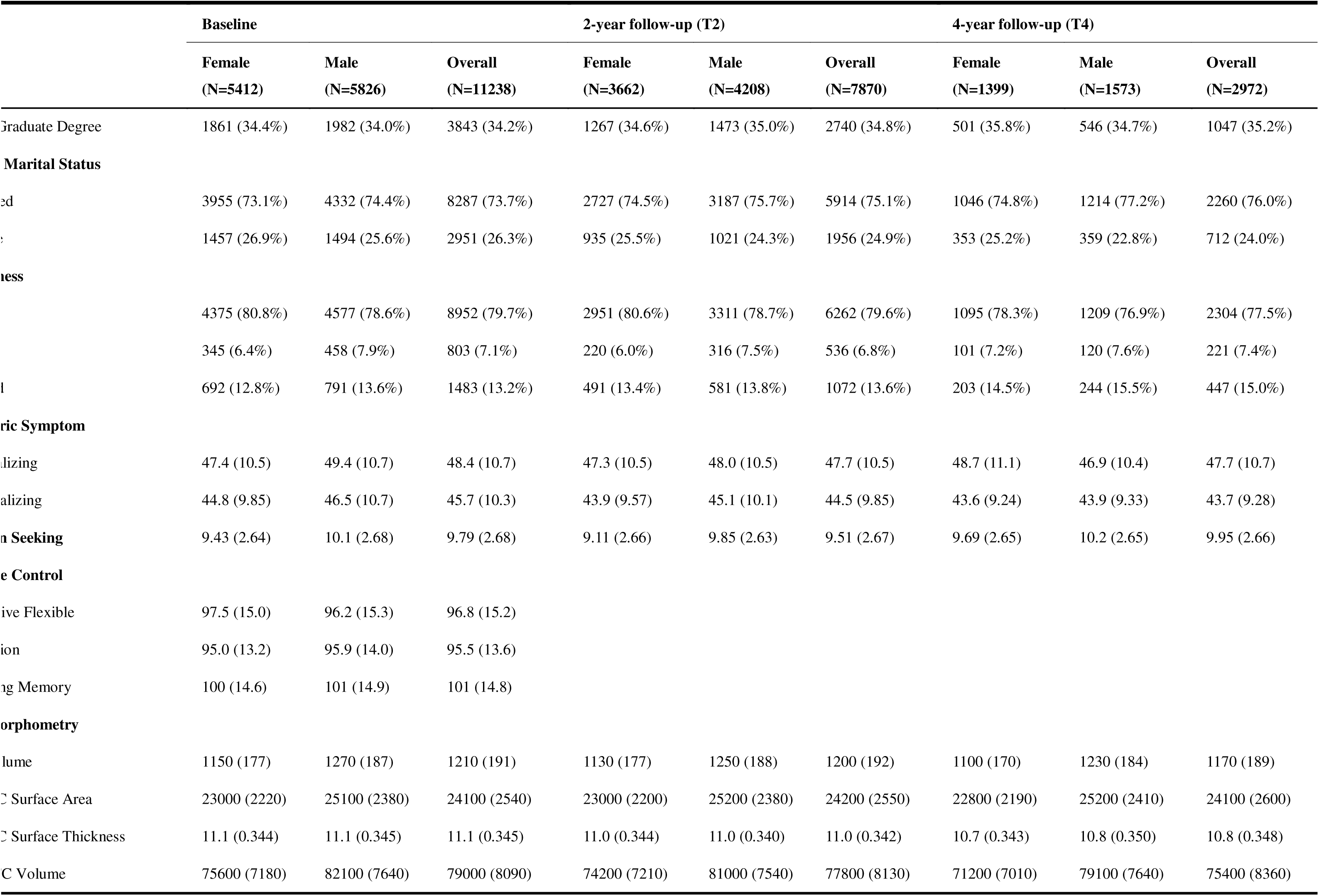
Descriptive Statistics of Demographic Characteristics of the ABCD study.

### Conceptualization of the Imbalance Score

The imbalance score quantified disparities between morphometric features of the VS and DLPFC —proxies for the socioemotional and cognitive-control systems, respectively. The VS, central to reward sensitivity, motivation salience and sensation-seeking, was selected to represent the socioemotional system. The DLPFC contributes to executive functions involved in top-down self-regulation, representing cognitive control (Figue1B, Supplement 6) (12,13).

The imbalance score was assessed along three dimensions: (1) absolute value, indicating the magnitude of disparity between dual systems; (2) direction of change, capturing developmental shifts (e.g., increasing vs. decreasing), which helps understand the dynamic interplay between the two systems; (3) growth rate, providing insight into the tempo of neurodevelopmental changes, with rapid growth indicating swift transitions and slower growth indicating gradual transitions.

### Neuroimaging Derivatives

Surface-based morphometry (SBM) was applied to structural MRI data to extract regional SA, ST and CV from the Desikan atlas. The SV of the VS was derived from ASEG parcellation. Detailed MRI acquisition and preprocessing protocols are available in prior publications (33). Ultimately, morphometric features included the SA, ST, CV for the DLPFC and SV for the VS.

### Psychopathology Measures

The Child Behavior Checklist (CBCL), a validated parent-reported tool for evaluating behavioral problems in youth aged 6 to 18, assessed psychopathological symptoms. Baseline t-scores from eight syndrome scales and two composite scores (internalizing and externalizing) were used. The internalizing domain encompasses anxious/depressed, withdrawn/depressed, and somatic complaints syndromes, while externalizing includes rule-breaking and aggressive behavior syndromes (35).

### Psychological Constructs of Dual Systems

Cognitive control and sensation-seeking were chosen as representative psychological constructs of the cognitive-control and socioemotional systems, respectively (Supplement 8).

**<u>Cognitive control</u>** is a multidimensional concept involving inhibition, working memory, and cognitive flexibility (36). To minimize systematic variance unrelated to cognitive control introduced by individual tasks (32,37,37), measures were quantified via latent variables extracted using principal component analysis (PCA) using age-corrected scores from NIH Toolbox neurocognitive tasks: the Flanker Task, List Sorting Working Memory Test, and Dimensional Change Card Sort Task (Supplement 1) (38).

**<u>Sensation-Seeking</u>** was evaluated using the sensation-seeking subscale of a modified version of the Urgency, Premeditation, Perseverance, and Sensation Seeking (UPPS-SS) Impulsive Behavior Scale for Children, which demonstrated high internal consistency and validity (39).

## Statistical Analysis

We first constructed the imbalance score and examined its reliability and validity. Initially, we analyzed the neural correlates of dual systems and associations with internalizing and externalizing dimensions. Given the widely accepted independence of cortical morphometric features (40,41), nested linear mixed models (LMMs) were used to examine the incremental validity of morphometric features (SA, CT, and CV) in predicting cognitive control, accounting for socio-demographic covariates, with site and family nested within site as random effects.

Marginal R-squared (ΔR^2^) values were computed to compare the relative importance of each feature (42). We then examined the relationships between morphometric features and psychopathology. The criterion-related validity of neural correlates with corresponding dual-systems psychological constructs was assessed using LMMs. The test-retest reliability of the imbalance score was evaluated using a mean-rating (k = 3), one-way random effects intra-class coefficient model (43) among participants with brain structural data available across all three waves.

Subsequently, we employed generalized additive models for location, scale, and shape (GAMLSS) to capture the developmental trajectory of the imbalance score using all available longitudinal data. GAMLSS is a flexible framework for modeling linear/non-linear growth trajectories and is recommended by the World Health Organization (44). The individual centile scores were computed to benchmark imbalance scores against normative age trends using a centile curve in GAMLSS (45). Centile scores quantified the deviations of the imbalance score relative to the normative distribution, where median values indicating relatively balanced development of the dual systems, and higher or lower values indicated a preference for the socioemotional or cognitive-control system, respectively. Details about model specification, stability and robustness are described in Supplement 4.

We derived the trajectory curve, variance, and growth rate of imbalance scores. The interpretation of the imbalance score should be additionally informed by how its individual components (DLPFC, VS) also change. Thus, separate trajectories for DLPFC and VS were also modeled using GAMLSS to capture regional developmental changes (Supplement 4). Furthermore, to preliminarily evaluate potential psychopathological changes across centile scores, we implemented a “tertile split” approach (46). Baseline participants were stratified into tertiles based on centile scores, and broad psychopathology scores were compared across the top, middle, and bottom 33% of centile scores using LMMs. To ensure equal statistical power across groups, propensity score matching was used to match groups demographics (47). Specifically, we assessed the fixed effects of tertile group membership and incorporated site and family structure as random effects.

Due to the non-linear nature of brain structural maturation, associations between imbalance scores and psychopathology dimensions (internalizing and externalizing) were evaluated using generalized additive models (GAMs) with penalized splines to capture non-linear effects. To account for the co-occurrence of internalizing and externalizing problems, internalizing scores were included as covariates when analyzing externalizing problems, and vice versa (48). We also assessed the interaction of sex and the smooth effect of age. Model selection was performed in a stepwise manner. These non-linear pattern relationships were further validated through LMMs and segmented polynomial models (Supplement 3). All results were considered significant at bilateral *p* < 0.05. All analyses were conducted in R 4.4.1 including *gamlss.dist* (49), *gamlss* (for GAMLSS) (45), *mgcv* (for GAM) (50) and *lmer4* (for LMM and segmented polynomial models) (51) packages.

## Sensitivity Analyses

To ensure the robustness of our findings, we conducted four sensitivity analyses. First, we tested the trajectory of the imbalance score and its associations with psychopathological symptoms using alternative imbalance score definitions based on the average deviation score (ADS) for SA, CT, and CV derived from a normative modeling framework (52) (Supplement 6.2 and Figure S3).

Normative deviation scores are considered more robust than raw brain features (53), as they account for deviations from healthy variation, reducing sample bias.

Second, we identified and removed outliers in brain morphology that produced extreme imbalance scores, potentially due to inaccurate segmentation or neurodevelopmental disorders (Supplement 6.1 and Figure S2).

Third, we compared the results of our imbalance scores with those from the competing indices to reduce the global effects and validate the unique effects of our selected proxies. We constructed a global imbalance score using global surface area (corresponding to DLPFC SA) and global subcortical volume (corresponding to VS SV) as proxies (Supplement 6.4) to alleviate potential confounding by global effects. Meanwhile, we calculated an alternative imbalance score using the difference between amygdala volume, a region also implicated in the Dual Systems Models and reward processing, and SA of DLPFC (Supplement 6.5) to validate the specificity of the imbalance score. This combined approach allowed us to assess whether the imbalance effects were unique to the VS-DLPFC axis or extended to other regions within the socioemotional and cognitive-control systems.

Finally, we tested the generalizability of our findings by replicating the analyses on the HCP-D dataset, hypothesizing that the imbalance scores would follow a similar developmental trajectory and exhibit comparable patterns in associations with psychopathology (Supplement 6.2 and Figure S2).

### Pre-Registration Statement

All aspects of our data analysis were pre-registered and made publicly available at https://osf.io/9bwfr/. Relevant components of the data analysis code can be accessed at https://github.com/MinfangKang773773/Dual-Systems-Imbalance.

### Covariates

Participant age, biological sex, parent-identified race, and parent-identified ethnicity were included as fixed-effect covariates in all LMMs and GAMs, along with parent-reported household income, parental highest education and marital status. In the GAMs, one sibling per family was randomly selected from baseline data to reduce model complexity. Collinearity was checked and confirmed by low Variance Inflation Factors (all < 10). Sex, race, ethnicity, and site were included in GAMLSS (Supplement 2).

## Results

### Neural Correlates of Dual Systems and Associations with Psychopathology

To determine the optimal neural correlate for the cognitive-control system, three surface morphological features were assessed by changes of marginal scores (ΔR_M_^2^) within nested LMMs. The SA (β =-0.093, SE = 0.10, 95%CI:-0.114 to-0.073, *p* < 0.001, ΔR_M_^2^ = 7.368 × 10^-3^) and cortical gray-matter volume (β =-0.077, SE = 0.10, 95%CI:-0.097 to-0.056, *p* < 0.001, ΔR_M_^2^ = 5.295×10^-3^) of the DLPFC were both significantly associated with cognitive control, whereas ST was not (β = 0.015, SE = 0.009, 95%CI:-0.003 to 0.034, *p* = 0.103, ΔR_M_^2^ = 9.103×10^-5^), as illustrated in Figure 2. SA exhibited the strongest predictive power for cognitive control and was therefore selected as the neural correlate for the cognitive-control system in subsequent analyses. Associations between single-system and psychopathology revealed distinct patterns (See Table 2). The broad internalizing symptoms were marginally associated with the socioemotional system (β =-1.819, SE = 0.98, *p* = 0.059) but not with the cognitive-control system. The broad externalizing symptoms were negatively associated with both the socioemotional system (β =-4.451, SE = 0.95, *p* < 0.001, ΔR_M_^2^ = 1.90×10^-3^) and cognitive-control system (β =-5.576, SE = 0.89, *p* < 0.001, ΔR_M_^2^ = 3.59×10^-3^), with stronger effects observed for the cognitive-control system.

**FIGURE 2.**
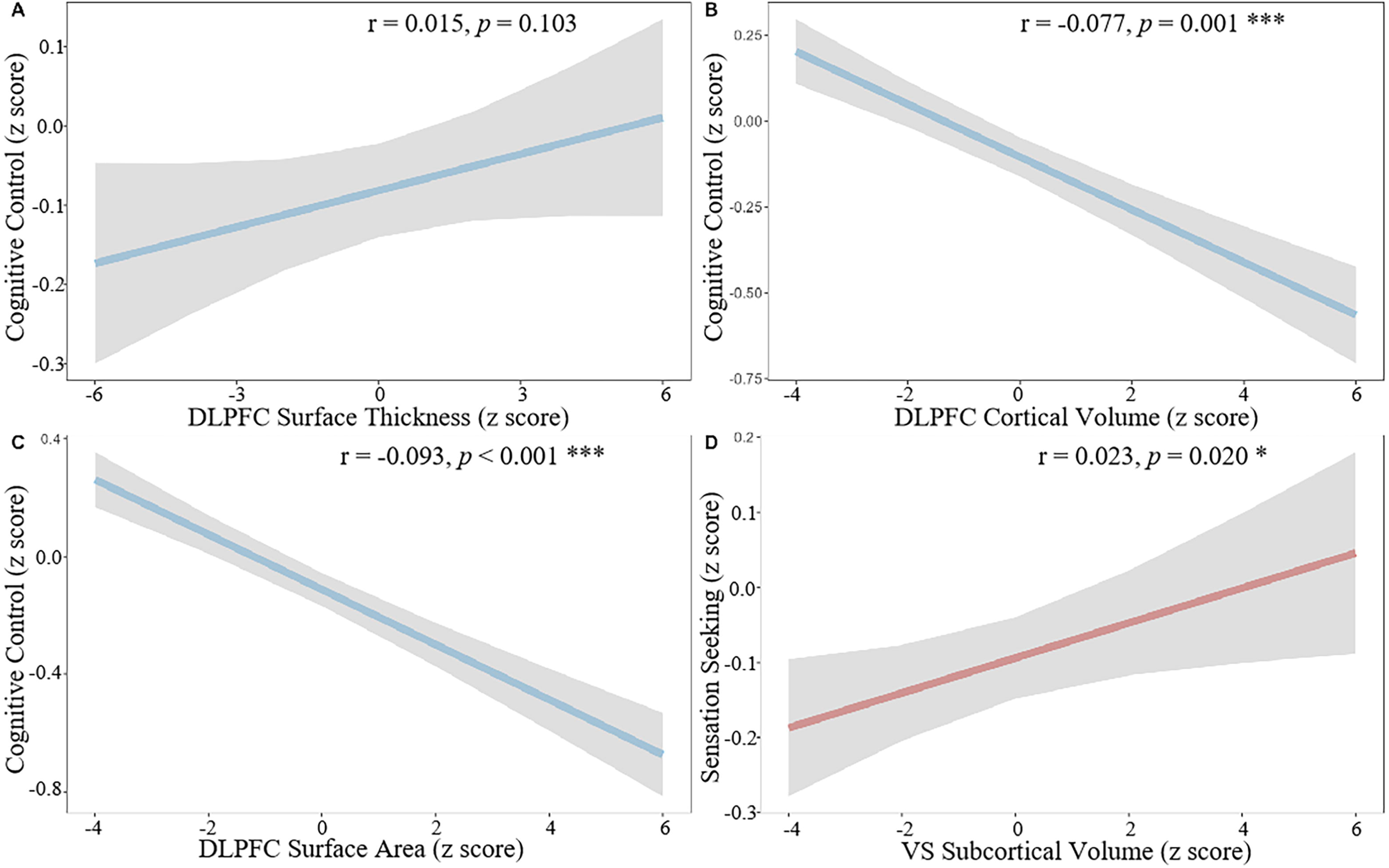
Associations between brain morphological features and psychological measurements of dual systems ***Panel A-D*** illustrate the associations between brain morphological features and psychological levels of Dual Systems Models using nested linear mixed models (LMMs). Dual Systems Models include the socioemotional system, involving the ventral striatum (VS) and psychological representation involving sensation-seeking, and the cognitive-control system, involving the dorsolateral prefrontal cortex (DLPFC) and psychological representation involving cognitive control. ***Panel A*** shows surface thickness (ST) of the dorsolateral prefrontal cortex (DLPFC) was not associated with cognitive control (β = 0.015, SE = 0.009, 95%CI:-0.003 to 0.034, *p* = 0.103, ΔR^2^ = 9.103×10^-5^). ***Panel B-C*** reveals the surface area (SA) (β =-0.093, SE = 0.10, 95%CI:-0.114 to-0.073, *p* < 0.001, ΔR^2^ == 7.368 × 10^-3^) and volume (β =-0.077, SE = 0.10, 95%CI:-0.097 to-0.056, *p* < 0.001, ΔR^2^ = 5.295×10^-3^) of DLPFC were associated with cognitive control. Marginal R²calculations suggest that the DLPFC SA may represent a primary neural correlate of the cognitive control system. ***Panel D*** shows the subcortical volume (SV) of the ventral striatum (VS) is significantly associated with sensation-seeking (β = 0.023, SE = 0.01, 95%CI: 0.0036 to 0.043, *p* = 0.020, ΔR^2^ = 4.636×10^-4^). * *P* < 0.05, ** *P* < 0.01, and *** *P* < 0.001.

**Table 2.** Associations between neural correlates of dual systems and psychopathology: evaluation and comparative importance.

|  | Socioemotional System (Subcortical Volume of VS) |  |  | Cognitive-control System (Surface Area of DLPFC) |  |  |
| --- | --- | --- | --- | --- | --- | --- |
|  | B(SE) | *p* | Δmarginal R<sup>2</sup> | B(SE) | *p* | Δmarginal R<sup>2</sup> |
| **Internalizing** |  |  |  |  |  |  |
| **Broadband internalizing** | -1.819(0.98) | 0.059<sup>+</sup> | **2.89×10<sup>-4</sup>** | -1.569(0.92) | 0.088 | 2.71×10<sup>-4</sup> |
| anxious/depressed | -1.043(0.56) | 0.064 | 3.09×10<sup>-4</sup> | -1.038(0.52) | 0.048* | **3.60×10<sup>-4</sup>** |
| withdrawn/depressed | -1.889(0.53) | < 0.001*** | **1.09×10<sup>-3</sup>** | -1.355(0.50) | 0.006** | 6.81×10<sup>-4</sup> |
| somatic complains | -1.074(0.56) | 0.051<sup>+</sup> | **3.05×10<sup>-4</sup>** | -0.120(0.53) | 0.821 | 5.27×10<sup>-6</sup> |
| **Externalizing** |  |  |  |  |  |  |
| **Broadband externalizing** | -4.451(0.95) | < 0.001*** | 1.89×10<sup>-3</sup> | -5.576(0.89) | < 0.001*** | **3.59×10<sup>-3</sup>** |
| rule-breaking behavior | -2.486(0.45) | < 0.001*** | 2.61×10<sup>-3</sup> | -2.507(0.42) | < 0.001*** | **3.24×10<sup>-3</sup>** |
| aggressive behavior | -2.315(0.51) | < 0.001*** | 1.79×10<sup>-3</sup> | -2.979(0.48) | < 0.001*** | **3.56×10<sup>-3</sup>** |
| **Other** |  |  |  |  |  |  |
| Social | -2.052(0.44) | < 0.001*** | 1.92×10<sup>-3</sup> | -1.906(0.41) | < 0.001*** | 1.95×10<sup>-3</sup> |
| Thought | -1.709(0.55) | 0.001** | 8.57×10<sup>-4</sup> | -1.467(0.51) | 0.004** | 7.61×10<sup>-4</sup> |
| Attention | -3.25(0.57) | < 0.001*** | 2.86×10<sup>-3</sup> | -3.600(0.53) | < 0.001*** | 4.15×10<sup>-3</sup> |

Abbreviations: VS, ventral striatum; DLPFC, dorsolateral prefrontal cortex.
$\Delta$ marginal $R^2$ the explanatory power of a single system, and bold font indicates the relatively more important predictors.
\* $P < 0.05$ , \*\* $P < 0.01$ , and \*\*\* $P < 0.001$ .

### Imbalance Score Generalizability and Validity

The personalized imbalance scores were calculated as the normalized difference between SV of VS minus SA of DLPFC. The imbalance scores demonstrated good test-retest reliability (ICC = 0.940, 95% CI: 0.936 to 0.944) (43). Structure-function analyses confirmed the validity of brain morphology features. As hypothesized, DLPFC SA statistically predicted cognitive control, and VS SV was associated with sensation-seeking (β = 0.023, SE = 0.01, 95%CI: 0.0036 to 0.043, *p* = 0.020) (Figure 2).

### Trajectory of the Dual-systems Imbalance

Using GAMLSS (45), we modeled non-linear age-related trends (in median and variance) stratified by sex, race and ethnicity, and accounting for site-specific ‘batch effects’ on structural MRI phenotypes in terms of multiple random effect parameters (44) in dual-systems imbalance. See Supplementary 4 for further details regarding GAMLSS model specification and estimation including model distribution (Supplementary 5.1), centile normalization (Supplementary 5.2), longitudinal centiles (Supplementary 5.3), model evaluation (Supplementary 5.4) and model sensitivity analyses (Supplementary 5.5).

The identified trajectory curve (Fig 3A) demonstrated a non-linear trajectory, remaining positive during early adolescence (ages 9–16), reflecting a greater prominence of the socioemotional system relative to the cognitive-control system. This persistent positive pattern suggests asynchronous and differential development between the two systems, potentially reflecting differential maturation rates in dual systems. Over time, scores decreased, reflecting a convergence between the two systems. This trend was accompanied by decreased variance and decelerating growth rates (Figure 3A-C). These changes implied that the cognitive-control system progressively caught up and lead to a more balanced interaction between dual systems.

**FIGURE 3.**
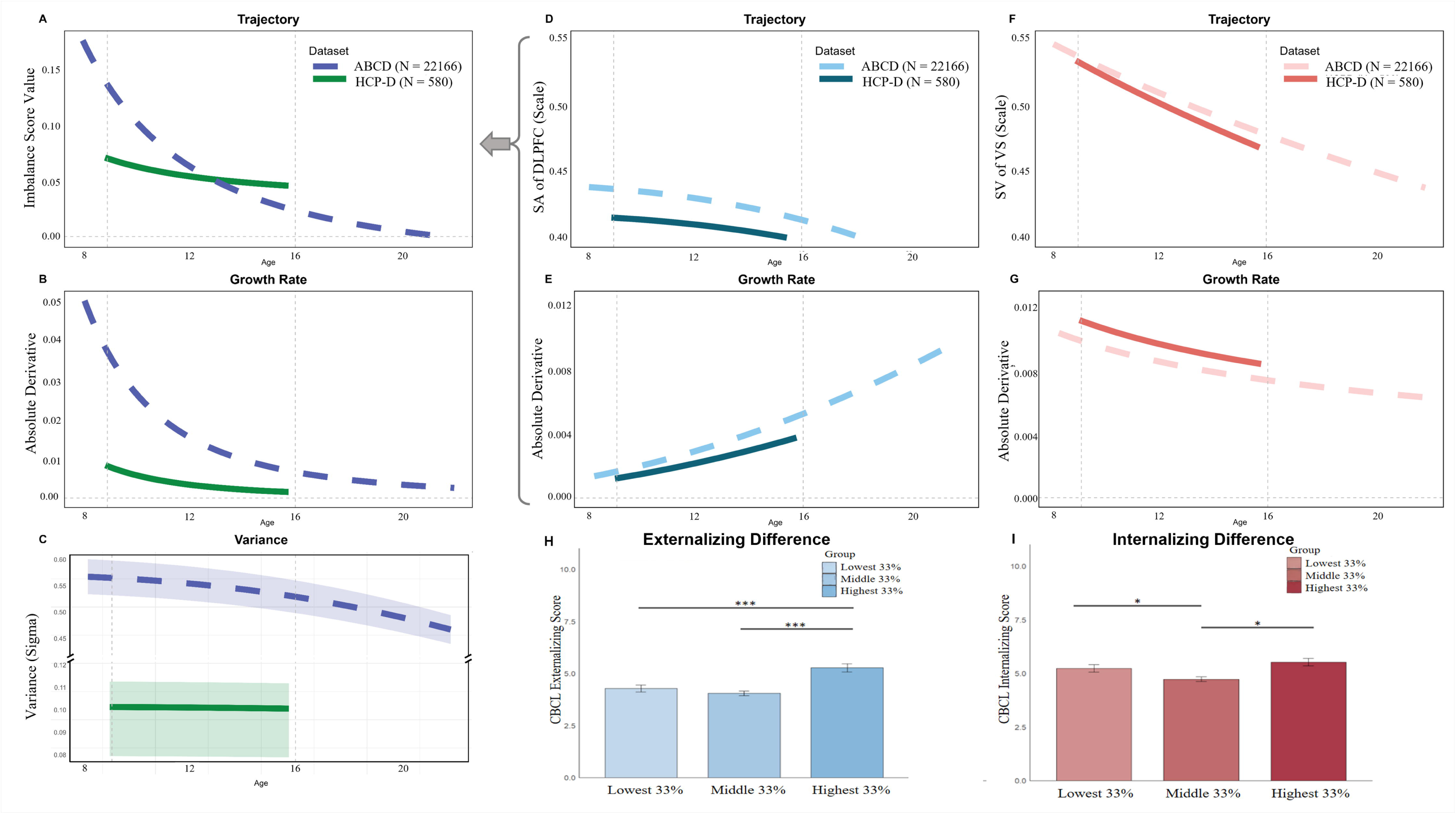
**Normative Trajectories and Variability of the Imbalance Scores Across Adolescence *Panels A-C*** depict the trajectory of the imbalance score analyzed in the Adolescent Brain Cognitive Development (ABCD) Study (age range: 9-16 years; N = 22166) and examined further in the Lifespan Human Connectome Project in Development (HCP-D) Study (age range: 8-21 years; N = 580). The normative trajectories are illustrated with generalized additive models for location, scale, and shape (GAMLSS), which adjusted for site-specific batch effects and controlled for covariates, including age, race, and ethnicity. The solid green lines represent results from the ABCD dataset, while the purple dotted lines represent findings from the HCP-D dataset. ***Panel A*** presents the normative trajectories of the imbalance scores, which were calculated as the subcortical volume (SV) of VS minus the surface area (SA) of DLPFC. The median imbalance scores demonstrate a non-linear developmental pattern, remaining positive and non-zero between ages 9 and 16 years in the ABCD dataset and ages 8 and 21 years in the HCP-D dataset. This consistently positive imbalance score reflects an asynchrony of the dual systems, with the socioemotional system exhibiting relative dominance over the cognitive-control system. The gradual decline in imbalance scores suggests that the disparity between the dual systems diminishes with age, reflecting a gradual convergence of the two systems as the cognitive-control system “catches up” to the socioemotional system. ***Panel B*** shows the rate of change in imbalance scores, calculated as the absolute value of the first derivative of the median trajectories. ***Panel C*** presents the trajectories of median between-subject variability and 95% confidence intervals for the imbalance scores. The variance of imbalance scores decreases during adolescence. ***Panels D-G*** depict the developmental trajectories of key brain regions representing the dual systems: the dorsolateral prefrontal cortex (DLPFC), associated with cognitive control, and the ventral striatum (VS), linked to socioemotional processing. As shown, the surface area of the DLPFC gradually decreased over time, with a relatively slow but gradually increasing change rate. The subcortical volume of the VS exhibits a faster rate of decrease during early adolescence, with the rate slowing as development progresses. Together, these trajectories explain the imbalance score’s developmental pattern, which is primarily driven by the relatively faster developmental rate and earlier maturity of the socioemotional system to the lagging cognitive-control system. ***Panels H-I*** shows the post-hoc analyses of LMMs for broad externalizing and internalizing scores. Linear mixed models (LMMs) with Tukey’s HSD post-hoc tests were conducted to explore group differences in externalizing and internalizing symptoms based on centile stratifications of imbalance scores. Groups were matched on demographics through propensity score matching (PSM), and each contained N = 3694 participants. * *P* < 0.05, ** *P* < 0.01, and *** *P* < 0.001.

**FIGURE 4.**
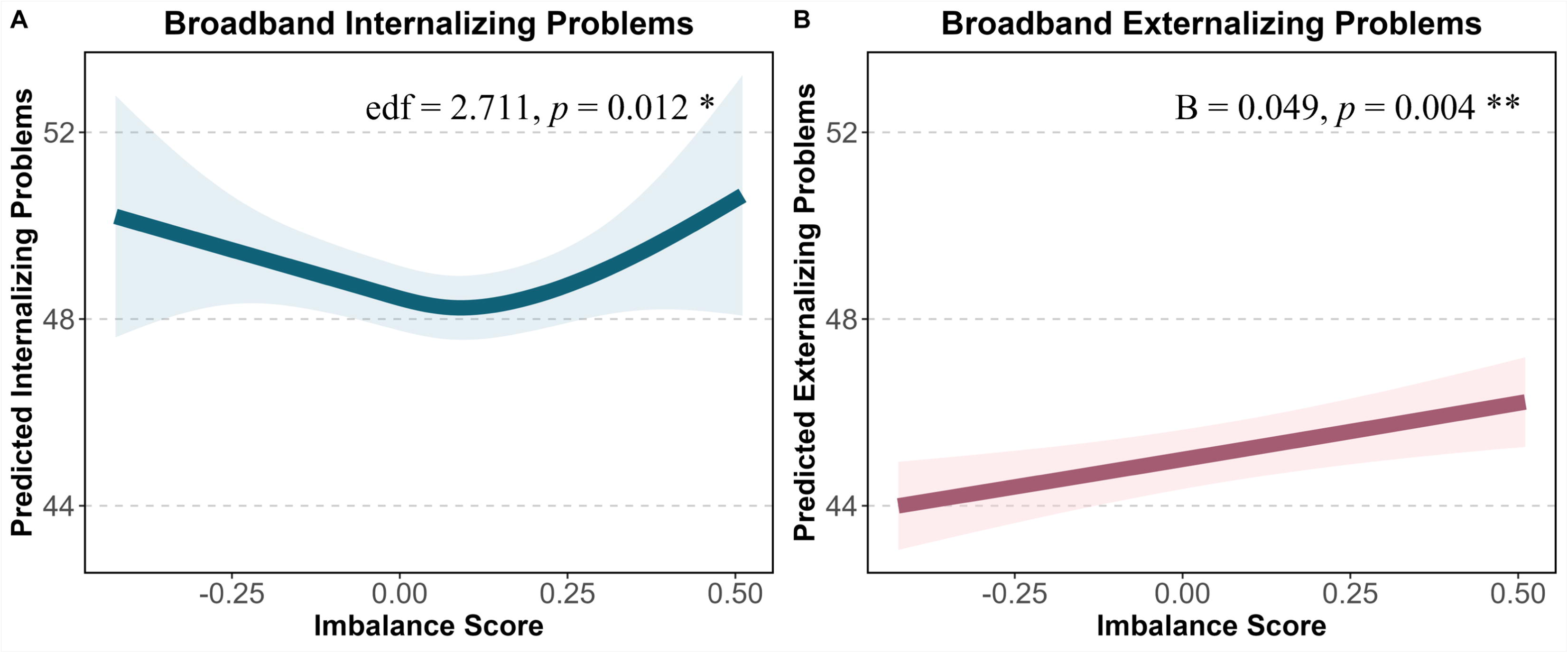
**Associations between the Imbalance Scores and Psychopathological Dimensions *Panels A-B*** illustrate the relationships between the imbalance scores and psychopathological dimensions using penalized splines within generalized additive models (GAMs). The models controlled for sex, age, race, ethnicity, parental education, family income, parental marital status, and other psychopathological dimensions. Fitted lines are the marginal relationship (female) pattern and shaded areas in both panels represent the 95% confidence intervals, derived from the standard errors of the predictions. ***Panel A*** demonstrates a significant non-linear association between the imbalance scores and broad internalizing symptoms (*edf* (imbalance score) = 2.711, *F* = 3.450, *p* = 0.012; *edf* (age)= 1.002, *F* = 8.601, *p* = 0.003) characterized by a similarly U-shaped curve with an inflection point at an imbalance score of approximately 0.10. ***Panel B*** shows a positive association with broad externalizing problems (B= 0.049, *SE* = 0.017, *p* = 0.004). These findings were further supported by linear mixed-effects models (LMMs) and segmented polynomial models (see Supplement 4). Abbreviations: edf, effective degrees of freedom. * *P* < 0.05, ** *P* < 0.01, and *** *P* < 0.001.

Investigation of the HCP-D dataset (ages 8-21) suggested similar trajectories (Figure 3A-C), and did outlier removal in ABCD (see Supplement 5).

Trajectories for individual systems revealed distinct trends (Figure 3D-G). DLPFC SA showed a stable, decreasing trend with an accelerating rate of change during adolescence. By contrast, VS SV decreased rapidly during early adolescence, with a decelerating rate of change. These were replicated in the HCP-D dataset, extending into early adulthood. These findings suggest that the imbalance trajectory was primarily driven by the socioemotional system’s faster maturation relative to the cognitive-control system.

Participants stratified into tertiles by imbalance centile scores exhibited significant differences in psychopathology (broad internalizing: *F*(_2, 10331_) = 3.438, *p* = 0.032; broad externalizing: *F*(_2, 10630_) = 10.68, *p* < 0.001). Participants in the middle group had significantly lower internalizing problem scores (B =-0.298, SE = 0.12, *p* = 0.015) compared to the upper-centile group. For externalizing scores, participants of both the lower group (B =-0.473, SE = 0.13, *p* < 0.001) and the middle group (B =-0.571, SE = 0.13, *p* < 0.001) had significantly lower scores compared to the upper-centile group (Figure 3H-I). Additionally, we conducted Tukey’s HSD post-hoc tests to explore pairwise differences between the tertile groups while controlling for multiple comparisons.

The results from both the mixed-effects models and post-hoc comparisons were interpreted in light of the random variability associated with site and family factors. There was no significant difference between the lower and middle-centile groups in post-hoc analyses.

### Imbalance Scores Specifically Statistically Predict Psychopathology Dimensions

We quantified the relationships between the imbalance score and internalizing/externalizing outcomes, separately. The LMMs results showed significant associations with externalizing symptoms (B = 1.793, SE = 0.730, *p* = 0.014) (See Supplementary 4.2). The GAMs (Table S3) revealed significant associations between imbalance scores and externalizing (B= 0.049, *SE* = 0.017, *p* = 0.004) and non-linear pattern of age (*edf* = 1.003, *F* = 13.2, *p* < 0.001), explaining 36.90% of the variance. The segmented polynomial models supported these relationships (see Supplementary 3 and Table S2).

Interestingly, a significant non-linear association with internalizing problems (*edf* (imbalance score) = 2.711, *F* = 3.450, *p* = 0.012; *edf* (age)= 1.002, *F* = 8.601, *p* = 0.003) (See Table S5) was identified, explaining 36.3% of the variance. Internalizing symptoms followed an approximately U-shaped trajectory, with both negative and positive imbalance extremes associated with higher symptoms. There was no significant interaction with sex. Segmented polynomial models supported the pattern (See Supplement 3 and Table S4).

For the competing indices, the global (B = 6.999, SE = 1.268, *p* < 0.001) and alternative imbalance (B = 2.898, SE = 0.752, *p* < 0.001) scores demonstrated robust associations with externalizing symptoms; however, their inconsistent and/or non-significant relationships with internalizing symptoms and reduced explanatory power compared to the original imbalance score (Supplement 6.4-6.5) suggested that the VS-DLPFC axis provided more precise and specific relationships transdiagnostically.

## Discussion

To enhance the precision of adolescent psychiatric prevention and reduce the burden of disorder onset, it is important to understand better individual-level neurobiological vulnerabilities to psychopathology. This study introduced a personalized imbalance score that quantified the developmental asynchrony between the cognitive-control and socioemotional systems using brain morphology metrics. Our findings support the Dual Systems Models by demonstrating that, during early adolescence, the dual systems exhibit asynchronous development, and the gap gradually decreased over time. Theoretically, the imbalance score reflects a relative measure of structural features associated with cognitive control and socioemotional processing, without implying that one is inherently superior or more mature. Rather, the dual systems contribute uniquely to behavior and psychopathology, and their balance or imbalance reflects the dynamic interplay rather than a simple hierarchy of “better” or “worse.” Additionally, we observed the decreasing variance and a slowing growth rate during adolescence, indicating decreased individual differences and a stabilization of the developmental trajectory over time. The imbalance score was positively associated with externalizing behaviors and showed an approximately U-shaped relationship with internalizing symptoms, reflecting distinct patterns of neurodevelopmental risk. These findings were independently supported in an independent sample from the HCP-D cohort, providing further support for the measure. Together, our results suggest that the imbalance score has potential clinical utility as a transdiagnostic predictor of both internalizing and externalizing psychopathology, offering valuable insights for targeted interventions and precision prevention.

Our findings highlight the relevance of structural brain metrics, consistent with studies showing that variations in cortical surface and thickness are reliably associated with neuropsychiatric concerns (54). Unlike functional connectivity studies relying on resting-state or task-based MRI (16) to investigate these theoretical hypotheses, we focused on structural data, which may be more genetically influenced, informative and predictive of psychopathology (40,55). Moreover, brain structure imprints on function, offering stable and reproducible findings (56). Dual Systems Models provide biologically plausible frameworks (10) for understanding these findings, supporting the theoretical rationale for the imbalance score.

In contrast to previous studies focusing on single behavioral tasks to operationalize the cognitive-control system, we adopted a broader latent approach. This method captures shared variance across core components and avoids the non-systematic, task-specific variance that may complicate interpretation (32). Furthermore, we examined multiple morphometric measures to understand the relationships between brain structure and psychopathological concerns (57,58).

Genetic architecture studies indicate that cortical surface area is more heritable and discoverable than thickness (28,29), aligning with our findings. The findings suggest the need for future studies to integrate multiple metrics to understand better brain-behavior relationships (29).

The personalized imbalance score may represent a novel neurobiological marker, collapsing two systems into a single variable that reflects variance in both. This measure may facilitate individualized understanding of neurodevelopmental trajectories, with implications for precision psychiatry. Meisel’s work on sensation-seeking and impulse control, which could be regarded as behavioral-level constructs, supports this approach, as the difference score predicted deviance (e.g., vandalism, substance use, school misconduct) (59), which aligns with our findings.

The observed asynchronous trajectory of imbalance can be traced to the specific systems involved. The gap is largely driven by the SA of the DLPFC, which shows a relatively slow but gradually increasing trend, and the SV of the VS, which exhibits a steeper decline. The prefrontal cortex undergoes synaptic pruning during adolescence, particularly in excitatory glutamate synapses, which enhances phasic dopamine signaling (60,61). This structural immaturity of the cognitive-control system may interact with elevated dopamine levels, potentially resulting in poor regulation of motivational signals. Adolescents exhibit heightened exploratory behaviors, novelty-seeking, and incentive salience, possibly reflecting dopamine’s role in promoting biologically salient incentives (62). These dynamics may explain adolescents’ cognitive immaturities, prefrontal regulation deficits, and heightened incentive-seeking during this developmental period.

Although the Dual Systems Models heuristically depict functional developmental trajectories, our study links these trajectories to structural maturation. Brain anatomy shapes and constrains neuronal activity (63), suggesting that structural and functional maturity largely align, although some divergence exists (64). The developmental changes implicated in the rate of change in imbalance scores may serve as a proxy for functional maturation within dual systems, potentially offering a measurable index for tracking neurodevelopment.

In accordance with predictions of Dual Systems frameworks (10,13,65), our results provide insight into Casey’s maturational imbalance hypothesis. The gradual decline in the imbalance score during adolescence reflects the cognitive-control system’s “catching up” with the socioemotional system. Specifically, VS volume decreased at a faster rate than DLPFC surface area increased, consistent with the systems’ converging strengths. These developmental changes align with theories proposing that synaptic pruning and myelination of prefrontal regions improve executive functioning, including response inhibition and risk-reward evaluation (65). However, due to the limited age range in our primary sample, we did not capture the theorized inflection point at which socioemotional system maturation plateaus. This plateau was possibly observed in our validation sample, where imbalance scores approached zero at ages 20–21 years.

Consistent with prior research, imbalance scores were associated with externalizing behaviors such as aggression and substance use (66,67), which have been linked to reduced prefrontal and subcortical volumes (68,69). Poor DLPFC function, coupled with heightened VS activity, may result in weak top-down regulation in the setting of strong motivational drives, thereby amplifying sensation-seeking. Conversely, the non-linear relationship between imbalance and internalizing symptoms suggests that low VS activity may impair hedonic processing, while diminished DLPFC function may exacerbate internalizing symptoms through poor behavioral flexibility.

While speculative, the findings and the larger literature suggest the potential involvement of moderating factors, consistent with the triadic model positing amygdala involvement in emotional regulation (70).

Given the frequent co-occurrence of internalizing and externalizing problems, these results suggest shared neurobiological mechanisms, akin to the p-factor in personality research.

Dysfunctions in fronto-striatal circuits have been implicated in multiple psychiatric disorders (71). Extending the imbalance score to a transdiagnostic framework could improve our understanding of comorbidities, a growing concern in mental health.

Several limitations should be noted. First, while we focused on key brain regions (DLPFC, VS) to construct the imbalance pattern, emerging evidence suggests that brain networks, rather than isolated regions of interest, may provide a more comprehensive understanding of neurodevelopment. Although achieving a one-to-one correspondence between neural markers and psychological states is ideal for improving inference precision (72), the VS and DLPFC are important regions in dual systems. Future research should integrate both spatial and temporal dynamics of brain networks (72). Second, our sample included participants aged 9 to 16 years, which does not encompass the full span of childhood/adolescence. Although the precise age ranges for childhood/adolescence vary in Dual Systems Models, our validation sample in the HCP-D (ages 8 to 21 years) aligns with Shulman’s framework (13), supporting the generalizability of our findings (Supplements 5). However, future studies should consider broader age ranges to capture developmental changes more comprehensively.

## Conclusions

We proposed and investigated a neuroanatomical imbalance score to quantify the divergent developmental trajectories of the socioemotional and cognitive-control systems based on the Dual Systems Models. This study is the first to reveal differential associations between the imbalance score and both internalizing and externalizing problems, suggesting that the imbalance score may serve as a transdiagnostic factor linked to developmental psychological concerns. Our findings suggest a personalized neurodevelopmental measure with transdiagnostic relevance, potentially offering novel insights into the etiology and precise prevention of psychopathology.

## Author Contributions

Dr. Zhang and BSc. Kang had full access to all of the data in the study and take responsibility for the integrity of the data and the accuracy of the data analysis.

Concept and design: J.T.Z, X.Y.F, M.F.K

Acquisition, analysis, or interpretation of data: All authors. Drafting of the manuscript: M.F.K, K.R.S, J.T.Z.

Critical revision of the manuscript for important intellectual content: Y.H.Z, X.Y.F, M.N.P. Statistical analysis: M.F.K, K.R.S, J.L.Z.

Obtained funding: J.T.Z, X.Y.F. Supervision: J.T.Z, X.Y.F.

## Data sharing

Data used in the preparation of this article were obtained from the Adolescent Brain Cognitive Development (ABCD) Study (https://abcdstudy.org), held in the NIMH Data Archive (NDA), and the Lifespan Human Connectome Project in Development (HCP-D) Study (https://humanconnectome.org/study/hcp-lifespan-development). ABCD is a multisite, longitudinal study designed to recruit more than 10,000 children age 9-10 and follow them over 10 years into early adulthood. The ABCD data repository grows and changes over time. The ABCD data used in this report came from NIMH Data Archive Digital Object Identifier (DOI: 10.15154/1520591). DOIs can be found at https://dx.doi.org/10.15154/1520591.

## Funding/Support

This study was supported by the National Natural Science Foundation of China [grant number 32371142 & 32171083, STI 2030 – Major Projects 2021ZD0200500], and 111 Project [BP0719032].

The ABCD Study® is supported by the National Institutes of Health and additional federal partners under award numbers U01DA041048, U01DA050989, U01DA051016, U01DA041022, U01DA051018, U01DA051037, U01DA050987, U01DA041174, U01DA041106, U01DA041117, U01DA041028, U01DA041134, U01DA050988, U01DA051039, U01DA041156, U01DA041025, U01DA041120, U01DA051038, U01DA041148, U01DA041093, U01DA041089, U24DA041123, U24DA041147. A full list of supporters is available at https://abcdstudy.org/federal-partners.html. A listing of participating sites and a complete listing of the study investigators can be found at https://abcdstudy.org/consortium_members/. ABCD consortium investigators designed and implemented the study and/or provided data but did not necessarily participate in the analysis or writing of this report.

The HCP-D study is supported by the National Institute of Mental Health of the National Institutes of Health under Award Number U01MH109589 and by funds provided by the McDonnell Center for Systems Neuroscience at Washington University in St. Louis.

This manuscript reflects the views of the authors and does not reflect the opinions or views of the NIH or ABCD consortium investigators.

The funders had no role in the design and conduct of the study; collection, management, analysis, and interpretation of the data; preparation, review, or approval of the manuscript; and decision to submit the manuscript for publication.

## Supplement Description

Supplement Methods, Results, Discussion, Figures S1-S5, Tables S1-S9

