## Supplementary material for "“Personalized Brain Morphometric Feature as a Transdiagnostic Predictor of Psychopathology: Insights from Dual Systems Models”": TABLE S1-TABLES9,FIGURE S1-FIGURE S5: Supplements_V8.0_FormatRevise.docx

**Supplemental Materials**

Title: “**Personalized Brain Morphometric Feature as a Transdiagnostic Predictor of Psychopathology: Insights for the Dual Systems Models**”

Authors: **Min-fang Kang^1^, Kun-ru Song^1^, Jia-lin Zhang^1^, Lin-xuan Xu^1^,**

**Ya-jie Zhang^1^, Zi-heng Zhao^1^, Hui-ying Deng^1^, Xiao-yi Fang^2*^, Marc N. Potenza^3,4,5,6,7,8^, Jin-tao Zhang^1*^**

**Corresponding authors:**

Xiao-Yi Fang, Ph.D., Professor

Institute of Developmental Psychology, Beijing Normal University, Beijing 100875, China

Beijing Normal University, No. 19, Xinjiekouwai street, Haidian District, Beijing 100875, China; Phone/Fax: +86 10 58808232,

Jin-Tao Zhang, Ph.D., Professor

State Key Laboratory of Cognitive Neuroscience and Learning and IDG/McGovern Institute for Brain Research, Beijing Normal University, No. 19, Xinjiekouwai street, Haidian District, Beijing 100875, China; Phone/Fax: +86 10 58800728,

**Content**

### TABLE S1. Selection Flow and Descriptive Statistics of Baseline Characteristics of the ABCD study

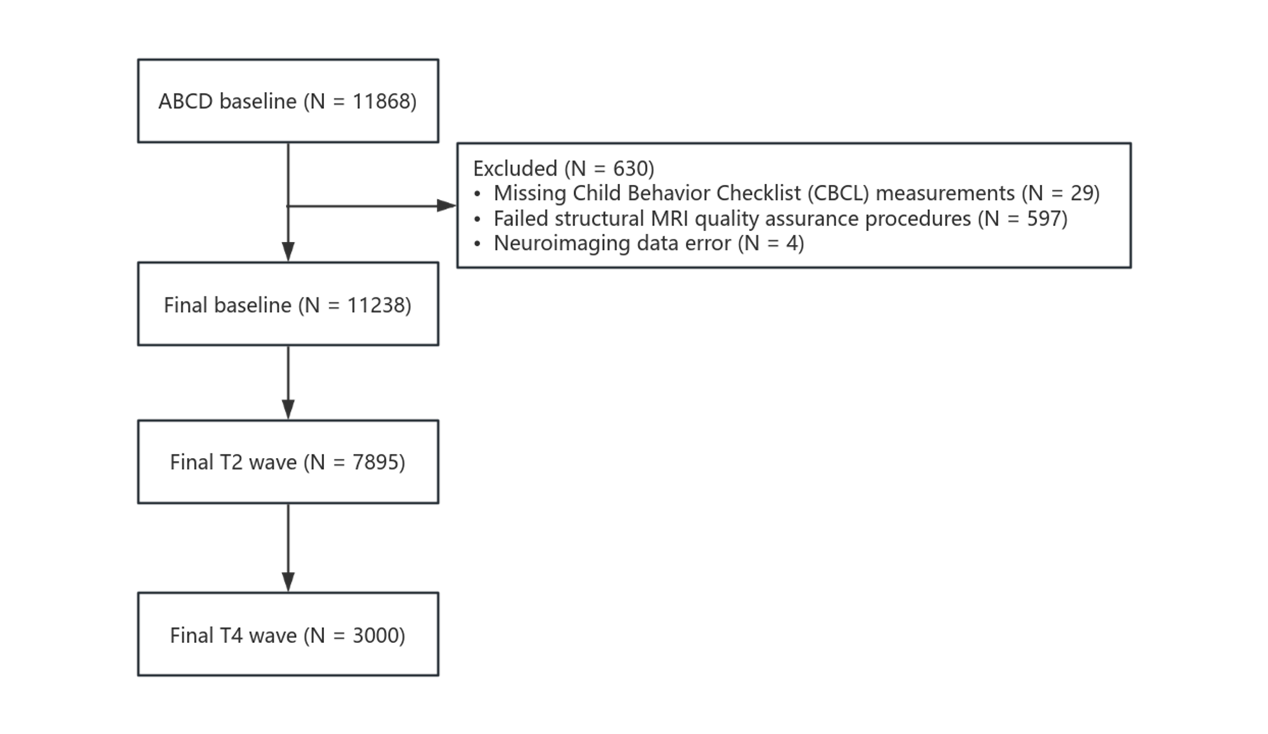

| Descriptive Statistics of Demographic Characteristics of the ABCD study | | | | | | | | | |
| --- | --- | --- | --- | --- | --- | --- | --- | --- | --- |
|  | **Baseline** | | | **2 years follow up (T2)** | | | **4 years follow up (T4)** | | |
|  | **Female (N=5412)** | **Male (N=5826)** | **Overall (N=11238)** | **Female (N=3662)** | **Male (N=4208)** | **Overall (N=7870)** | **Female (N=1399)** | **Male (N=1573)** | **Overall (N=2972)** |
| **Age (years)** | 9.90 (0.623) | 9.94 (0.627) | 9.92 (0.625) | 11.9 (0.651) | 12.0 (0.651) | 12.0 (0.652) | 14.1 (0.687) | 14.1 (0.698) | 14.1 (0.693) |
| **Race** |  |  |  |  |  |  |  |  |  |
| White | 3438 (63.5%) | 3821 (65.6%) | 7259 (64.6%) | 2369 (64.7%) | 2875 (68.3%) | 5244 (66.6%) | 934 (66.8%) | 1131 (71.9%) | 2065 (69.5%) |
| Black | 871 (16.1%) | 864 (14.8%) | 1735 (15.4%) | 540 (14.7%) | 565 (13.4%) | 1105 (14.0%) | 176 (12.6%) | 154 (9.8%) | 330 (11.1%) |
| Asian | 135 (2.5%) | 122 (2.1%) | 257 (2.3%) | 81 (2.2%) | 79 (1.9%) | 160 (2.0%) | 34 (2.4%) | 37 (2.4%) | 71 (2.4%) |
| American Indian/Alaska Native | 32 (0.6%) | 24 (0.4%) | 56 (0.5%) | 25 (0.7%) | 19 (0.5%) | 44 (0.6%) | 15 (1.1%) | 10 (0.6%) | 25 (0.8%) |
| Native Hawaiian/Other Pacific | 6 (0.1%) | 9 (0.2%) | 15 (0.1%) | 6 (0.2%) | 7 (0.2%) | 13 (0.2%) | 2 (0.1%) | 2 (0.1%) | 4 (0.1%) |
| Other | 667 (12.3%) | 688 (11.8%) | 1355 (12.1%) | 475 (13.0%) | 476 (11.3%) | 951 (12.1%) | 176 (12.6%) | 170 (10.8%) | 346 (11.6%) |
| Mixed | 263 (4.9%) | 298 (5.1%) | 561 (5.0%) | 166 (4.5%) | 187 (4.4%) | 353 (4.5%) | 62 (4.4%) | 69 (4.4%) | 131 (4.4%) |
| **Ethnicity** |  |  |  |  |  |  |  |  |  |
| Hispanic/Latino/Latina | 4308 (79.6%) | 4641 (79.7%) | 8949 (79.6%) | 2939 (80.3%) | 3424 (81.4%) | 6363 (80.9%) | 1121 (80.1%) | 1254 (79.7%) | 2375 (79.9%) |
| Non-Hispanic/Latino/Latina | 1104 (20.4%) | 1185 (20.3%) | 2289 (20.4%) | 723 (19.7%) | 784 (18.6%) | 1507 (19.1%) | 278 (19.9%) | 319 (20.3%) | 597 (20.1%) |
| **Household Income** |  |  |  |  |  |  |  |  |  |
| <$50k | 1570 (29.0%) | 1607 (27.6%) | 3177 (28.3%) | 1023 (27.9%) | 1105 (26.3%) | 2128 (27.0%) | 380 (27.2%) | 396 (25.2%) | 776 (26.1%) |
| $50k-$100k | 1413 (26.1%) | 1516 (26.0%) | 2929 (26.1%) | 1035 (28.3%) | 1144 (27.2%) | 2179 (27.7%) | 400 (28.6%) | 448 (28.5%) | 848 (28.5%) |
| >$100k | 2429 (44.9%) | 2703 (46.4%) | 5132 (45.7%) | 1604 (43.8%) | 1959 (46.6%) | 3563 (45.3%) | 619 (44.2%) | 729 (46.3%) | 1348 (45.4%) |
| **Parental Education** |  |  |  |  |  |  |  |  |  |
| < High School Diploma | 654 (12.1%) | 660 (11.3%) | 1314 (11.7%) | 401 (11.0%) | 434 (10.3%) | 835 (10.6%) | 130 (9.3%) | 147 (9.3%) | 277 (9.3%) |
| High School Diploma/GED | 823 (15.2%) | 915 (15.7%) | 1738 (15.5%) | 562 (15.3%) | 633 (15.0%) | 1195 (15.2%) | 200 (14.3%) | 234 (14.9%) | 434 (14.6%) |
| Some College | 714 (13.2%) | 768 (13.2%) | 1482 (13.2%) | 483 (13.2%) | 534 (12.7%) | 1017 (12.9%) | 188 (13.4%) | 204 (13.0%) | 392 (13.2%) |
| Bachelor | 1360 (25.1%) | 1501 (25.8%) | 2861 (25.5%) | 949 (25.9%) | 1134 (26.9%) | 2083 (26.5%) | 380 (27.2%) | 442 (28.1%) | 822 (27.7%) |
| Post-Graduate Degree | 1861 (34.4%) | 1982 (34.0%) | 3843 (34.2%) | 1267 (34.6%) | 1473 (35.0%) | 2740 (34.8%) | 501 (35.8%) | 546 (34.7%) | 1047 (35.2%) |
| **Parental Marital Status** |  |  |  |  |  |  |  |  |  |
| Married | 3955 (73.1%) | 4332 (74.4%) | 8287 (73.7%) | 2727 (74.5%) | 3187 (75.7%) | 5914 (75.1%) | 1046 (74.8%) | 1214 (77.2%) | 2260 (76.0%) |
| Single | 1457 (26.9%) | 1494 (25.6%) | 2951 (26.3%) | 935 (25.5%) | 1021 (24.3%) | 1956 (24.9%) | 353 (25.2%) | 359 (22.8%) | 712 (24.0%) |
| **Handedness** |  |  |  |  |  |  |  |  |  |
| Right | 4375 (80.8%) | 4577 (78.6%) | 8952 (79.7%) | 2951 (80.6%) | 3311 (78.7%) | 6262 (79.6%) | 1095 (78.3%) | 1209 (76.9%) | 2304 (77.5%) |
| Left | 345 (6.4%) | 458 (7.9%) | 803 (7.1%) | 220 (6.0%) | 316 (7.5%) | 536 (6.8%) | 101 (7.2%) | 120 (7.6%) | 221 (7.4%) |
| Mixed | 692 (12.8%) | 791 (13.6%) | 1483 (13.2%) | 491 (13.4%) | 581 (13.8%) | 1072 (13.6%) | 203 (14.5%) | 244 (15.5%) | 447 (15.0%) |
| **Psychiatric Symptom** |  |  |  |  |  |  |  |  |  |
| Internalizing | 47.4 (10.5) | 49.4 (10.7) | 48.4 (10.7) | 47.3 (10.5) | 48.0 (10.5) | 47.7 (10.5) | 48.7 (11.1) | 46.9 (10.4) | 47.7 (10.7) |
| Externalizing | 44.8 (9.85) | 46.5 (10.7) | 45.7 (10.3) | 43.9 (9.57) | 45.1 (10.1) | 44.5 (9.85) | 43.6 (9.24) | 43.9 (9.33) | 43.7 (9.28) |
| **Sensation Seeking** | 9.43 (2.64) | 10.1 (2.68) | 9.79 (2.68) | 9.11 (2.66) | 9.85 (2.63) | 9.51 (2.67) | 9.69 (2.65) | 10.2 (2.65) | 9.95 (2.66) |
| **Cognitive Control** |  |  |  |  |  |  |  |  |  |
| Cognitive Flexible | 97.5 (15.0) | 96.2 (15.3) | 96.8 (15.2) |  |  |  |  |  |  |
| Inhibition | 95.0 (13.2) | 95.9 (14.0) | 95.5 (13.6) |  |  |  |  |  |  |
| Working Memory | 100 (14.6) | 101 (14.9) | 101 (14.8) |  |  |  |  |  |  |
| **Brain Morphometry** |  |  |  |  |  |  |  |  |  |
| VS Volume | 1150 (177) | 1270 (187) | 1210 (191) | 1130 (177) | 1250 (188) | 1200 (192) | 1100 (170) | 1230 (184) | 1170 (189) |
| DLPFC Surface Area | 23000 (2220) | 25100 (2380) | 24100 (2540) | 23000 (2200) | 25200 (2380) | 24200 (2550) | 22800 (2190) | 25200 (2410) | 24100 (2600) |
| DLPFC Surface Thickness | 11.1 (0.344) | 11.1 (0.345) | 11.1 (0.345) | 11.0 (0.344) | 11.0 (0.340) | 11.0 (0.342) | 10.7 (0.343) | 10.8 (0.350) | 10.8 (0.348) |
| DLPFC Volume | 75600 (7180) | 82100 (7640) | 79000 (8090) | 74200 (7210) | 81000 (7540) | 77800 (8130) | 71200 (7010) | 79100 (7640) | 75400 (8360) |

Upper is the CONSORT flow diagram of participants in the ABCD study.

### Supplement 1. Principal component analysis (PCA) of cognitive control

Cognitive control refers to a set of top-down mental processes that enable goal-directed behavior, particularly when automatic or instinctual actions are insufficient or inappropriate (1). Three core components of cognitive control have been proposed: inhibition, working memory (WM), and cognitive flexibility, representing elements of executive functioning (2).

In this study, we assessed these components using three tasks from the NIH Toolbox cognition battery: the Flanker Inhibitory Control and Attention Test (inhibition), the List Sorting Working Memory Test (working memory), and the Dimensional Change Card Sort Test (cognitive flexibility) (3). We performed principal component analysis (PCA) to extract a latent cognitive control factor, regressing task performance onto a single latent variable (4). Bartlett’s test (5) of sphericity was significant, $\chi^{2}\left( 2 \right)=140.05, p<0.001$, and an overall measure of sampling adequacy (6) was 0.6, exceeding the minimum recommended threshold of 0.50, suggesting that the data were appropriate for factor analysis. All analyses used standardized scores, and age-related differences were corrected.

The PCA revealed one principal component with the highest eigenvalue accounting for 54.33% of the total variance, which we retained as the cognitive-control factor. Factor loadings for the three tasks on this first principal component were as follows: inhibition = 0.61, cognitive flexibility = 0.62, and working memory = 0.49, all in the moderate range. This principal component was subsequently used as the composite cognitive control score for subsequent analyses.

### Supplement 2. Covariates

As in previous studies utilizing the ABCD data (7–10), we have included a set of demographics, including youth’s age, biological sex, parent-identified race, and parent-identified ethnicity as covariates in linear mixed models (LMMs). Multiple caretaker measures associate with youth mental health and development, including parental education associating with externalizing and total problems (9) and family income and financial adversity account for unique variance in reports of children mental health (9). For the GAMLSS analysis, we included sex, race and ethnicity as covariates, as in previous studies (11). The same covariates were included in the analyses of the HCP-D sample.

### Supplement 3. Generalized Additive Model: Association between Imbalance Score with Psychopathological Dimensions

Given the non-linear nature of brain structural maturation, we applied penalized splines in generalized additive models (GAMs) to examine both linear and non-linear associations between the imbalance scores and broad internalizing/externalizing symptoms. Covariates included sex, age, race, ethnicity, parental education, familial income, and parental marital status, with site modeled as a random effect. To simplify the analysis, one sibling per family was randomly selected from baseline data. Due to the high correlation and comorbidity between internalizing and externalizing problems, internalizing symptoms were included as covariates when examining externalizing problems, and vice versa. We initially employed linear mixed models (LMMs) and segmented polynomial models to explore the relationships patterns, then used GAMs to capture the precise relationships. Model selection was based on chi-square tests, and smoothing parameters were determined using the generalized cross-validation (GCV) criterion (12).

Based on prior results, socioemotional system was more informative for predicting internalizing dimensions (e.g., withdrawn/depressed and somatic complaints), while the cognitive-control system was more critical for predicting externalizing dimensions (e.g., rule-breaking behavior and aggressive behavior); see Table 1.

#### 4.1 Associations between Imbalance Score and Broad Internalizing Concerns

The LMMs indicated that the imbalance score was not a significant linear predictor of the broad internalizing problems (B = -0.126, SE = 0.759, *p* = 0.097). However, residual analysis suggested a non-linear cubic relationship, indicating that linear models did not adequately capture the underlying data structure. Further exploration with segmented polynomial models, testing 2-5 terms for the imbalance score, revealed a significant quadratic effect shown in Table S4, supporting the presence of a non-linear relationship.

**Table S4. The relationships between the broad internalizing and the imbalance scores in segmented polynomial models.**

| **Segmented Polynomial models** | **Imbalance score, 1** | **Imbalance score, 2** | **Imbalance score, 3** | **Imbalance score, 4** | **Imbalance score, 5** |
| --- | --- | --- | --- | --- | --- |
| 2 | B = -14.79, SE = 9.060, *p* = 0.103 | **B = 23.79, SE = 8.574, *p* = 0.005**** | - | - | - |
| 3 | B = -14.88, SE = 9.060, *p* = 0.101 | **B = 23.80, SE = 8.574, *p* = 0.005**** | B = 8.281, SE = 8.493, *p* = 0.329 | - | - |
| 4 | B = -14.88, SE = 9.060, *p* = 0.100 | **B = 23.75, SE = 8.575, *p* = 0.005**** | B = 8.282, SE = 8.493, *p* = 0.329 | B = 7.428, SE = 8.490, *p* = 0.381 | - |
| 5 | B = -14.88, SE = 9.061, *p* = 0.100 | **B = 23.75, SE = 8.575, *p* = 0.005**** | B = 8.281, SE = 8.493, *p* = 0.329 | B = 7.430, SE = 8.491, *p* = 0.381 | B = 0.548, SE = 8.497, *p* = 0.948 |

GAMs were then fitted using N = 9,373 participants (after random sibling selection). Familial effects were confirmed through model contrasts, revealing that eliminating the familial effect increased model explanatory power. Six GAMs were fitted and compared using chi-square tests. The reduced model, with non-smooth terms and a random site effect which was specified by adding the terms *s(site, bs="re")* to the model formula (12), revealed that the imbalance score was not significant (B = -1.447, SE = 0.802, *p* = 0.071) and the site effect was marginally significant (*edf* = 0.729, *F* = 2.692, *p* = 0.054). The second model added smooth effect to imbalance score and the smooth term for the imbalance score was significant (*edf* = 2.778, *F* = 3.701, *p* = 0.008$, GCV=73.117$), and the model comparison showed that including the smooth term improved the fit (*F*(2.538, 9359) = 4.839, *p* = 0.004). The third model replaced the random effect of site within a fixed effect to prevent overfitting and showed model fit significantly better (*F* (20.043, 9339.5) = 4.719, *p* < 0.001). To address heteroscedasticity observed in residual distributions, a gamma distribution with a log link function was applied in the fourth model, which improved residual normality and QQ plot fit (*edf* = 2.708, *F* = 3.451, *p* = 0.011).This model was preferred based on both GCV ($GCV=0.031$). The fifth and sixth models included smooth terms for age and an interaction between imbalance score and sex, respectively. The smooth term of age-term was significant (*edf* = 2.711, *F* = 3.450, *p* = 0.012, $GCV=0.031$), and the model fitted better (*F* (0.007, 9340) = 0.767, *p* = 0.019), while the sex interaction was not significant (Male*: edf =* 0.015*, F* = 0.225, *p* = 0.936; Female: *edf =* 1.001*, F* = 1.688, *p* = 0.194). We finally utilized the fifth model as the quantified pattern, and we selected the smoothing parameter, including knot and penalty, by GCV criterion. The final model was:

$$\log\left( CBCL_{Internal},i \right)=a_{0}+\alpha_{1}cr\left( imb_{i} \right)+\alpha_{2}cr\left( age_{i} \right)+\alpha_{3}race_{i}+ \alpha_{4}ethnicity_{i}+\alpha_{5}income_{i}+\alpha_{6}education_{i}+\alpha_{7}marital_{i}+\alpha_{8}site_{i}+ \varepsilon_{i}, CBCL_{internalizing} \sim Gammagamma=1，CBCL\_Internal \sim Gamma$$

The model explained 35.2% of the variance (GCV = 0.031), with estimated degrees of freedom (*edf*) for imbalance score and age being 2.486 and 1.002, respectively.

We utilized generalized cross validation (GCV) to validate our model. Validation using the GCV criterion and the *gam.check* routine confirmed successful convergence (k-index(the k-index is the ratio of the model estimated scale parameter to an estimate based on differencing neighboring residuals) = 1, p(the p-value is for the residual randomization test) > 0.05), indicating good model fit.

**Table S5. The model selections in the relationships with the broad internalizing problems in GAMs.**

| **GAMs** | **Term: Imbalance score** | **Term: Age** | **Term: Sex (male)** | **Term: Imbalance score * sex** | **Term: Site** | **Model Comparison** |
| --- | --- | --- | --- | --- | --- | --- |
| Reduced model | B = -1.447, SE = 0.802,  *p* = 0.071. | **B = 0.364, SE = 0.143,**  ***p* < 0.011*** | **B = 0.829, SE = 0.180,**  ***p* < 0.001***** | - | *edf* = 0.729, *F* = 2.692,  *p* = 0.054. | - |
| Model 2 | ***edf* = 2.778, *F* = 3.701,**  ***p* = 0.008**** | **B = 0.365, SE = 0.143,**  ***p* = 0.011*** | **B = 0.814, SE = 0.180,**  ***p* < 0.001***** | - | *edf* = 0.740, *F* = 2.830,  *p* = 0.051. | ***F* (2.538, 9359) = 4.839,**  ***p =* 0.004**** |
| Model 3 | ***edf* = 2.757, *F* = 3.312,**  ***p* = 0.014*** | **B = 0.449, SE = 0.144,**  ***p* = 0.002**** | **B = 0.839, SE = 0.179,**  ***p* < 0.001***** | - |  | ***F* (20.043,9339.5) = 4.719, *p <* 0.001***** |
| Model 4 | ***edf =* 2.708*, F* = 3.451,**  ***p* = 0.011∗** | **B = 0.008, SE = 0.003,**  ***p* = 0.003**** | **B = 0.018, SE = 0.004,**  ***p* < 0.001***** | - |  | **-** |
| Model 5 | ***edf =* 2.711*, F* = 3.450,**  ***p* = 0.012∗** | ***edf =* 1.002*, F* = 8.601,**  ***p* = 0.003**** | **B = 0.018, SE = 0.004,**  ***p* < 0.001***** | - |  | ***F* (0.007,9340) = 0.767,**  ***p =* 0.019*** |
| Model 6 | ***edf =* 2.744*, F* = 3.486,**  ***p* = 0.011*** | ***edf =* 1.000*, F* = 8.690,**  ***p* = 0.003**** | **-** | Male*: edf =* 0.015*, F* = 0.225,  *p* = 0.936  Female: *edf =* 1.001*, F* = 1.688,  *p* = 0.194 |  | *F* (1.073,9338.5) = 1.937,  *p =* 0.163 |

#### 4.2 Association between Imbalance Scores and Broad Externalizing Concerns

LMMs revealed a significant association between the imbalance score and broad externalizing problems (B = 1.793, SE = 0.730, *p* = 0.014). This effect persisted across polynomial models segmented by 2-5 terms (See Table-S2), although residual analysis indicated potential non-linearity in the relationship.

**Table S2. The relationships between the broad externalizing and the imbalance scores in segmented polynomial models.**

| **Segmented Polynomial models** | **Imbalance score, 1** | **Imbalance score, 2** | **Imbalance score, 3** | **Imbalance score, 4** | **Imbalance score, 5** |
| --- | --- | --- | --- | --- | --- |
| 2 | **B = 21.25, SE = 8.714, *p* = 0.014*** | **B = -13.74, SE = 8.274, *p* = 0.097.** | - | - | - |
| 3 | **B = 21.30, SE = 8.714, *p* = 0.014*** | **B = -13.75, SE = 8.274, *p* = 0.097.** | B = -6.146, SE = 8.200, *p* = 0.453 | - | - |
| 4 | **B = 21.30, SE = 8.714, *p* = 0.014*** | **B = -13.75, SE = 8.275, *p* = 0.097.** | B = -6.146, SE = 8.200, *p* = 0.453 | B = -0.392, SE = 8.197, *p* = 0.962 | - |
| 5 | **B = 21.27, SE = 8.713, *p* = 0.014*** | **B = -13.73, SE = 8.274, *p* = 0.097.** | B = -6.174, SE = 8.199, *p* = 0.451 | B = 0.434, SE = 8.197, *p* = 0.957 | **B = 13.71, SE = 8.201, *p* = 0.095.** |

We then fitted six GAMs, with the reduced model containing a linear effect for imbalance score and a random effect for site, as well as covariates in linear pattern. The imbalance score was significant (B = 1.800, SE = 0.768, *p* = 0.019, GCV = 67.339), while the smooth term for site was not (*edf* = 0.003, *F* = 0.003, *p* = 0.316). In the second model, adding a smooth term for the imbalance score improved model fit slightly (*edf* = 2.851, *F* = 2.909, *p* = 0.025), and there was no significant positive improvement of the model fit ( *F* (2.696, 9363) = 2.575, *p =* 0.058), which demonstrated the smooth effect would add complexity but no considerable model fit improvement. Adjusting site to a fixed effect in the third model maintained the significant association between imbalance scores and externalizing concerns (*B* = 2.190, *SE* = 0.799, *p* = 0.006$, GCV=67.126$), with a significantly better fit compared to the second model (*F* (20.994, 9363) = 3.418, *p <* 0.001). Meanwhile, we applied the gamma distribution and the log link function to address heteroscedasticity, which significantly improved model performance (*B =* 0.049*, SE* = 0.017, *p* = 0.004, GCV = 0.031). The fifth and sixth models examined the smooth term of age and interaction with sex, respectively, showing that the fifth model fit well (*B* = 0.049*, SE* = 0.017, *p* = 0.004; *edf* (age) *=* 1.003*, F* = 13.2, *p* < 0.001; $GCV=0.031)$ and significantly fitted better (*F* (0.006, 9342) = 0.289, *p =* 0.018), and the sixth model showed that the interaction with sex was not significant improve the model fit (Male*: edf =* 1.001*, F* = 1.061, *p* = 0.303; Female: *edf =* 2.106*, F* = 4.711, *p* = 0.003; *F* (2.723, 9342) = 2.680, *p* = 0.051). The whole results were shown in Table-S3.

We finally used the fifth model as our final relationship pattern, and the imbalance scores were significantly associated with broad externalizing concerns and the final model was:

$$\log\left( CBCL_{External},i \right)=a_{0}+\alpha_{1}imb_{i}+\alpha_{2}cr\left( age_{i} \right)+ \alpha_{3}ethnicity_{i}+\alpha_{4}income_{i}+\alpha_{5}education_{i}+\alpha_{6}marital_{i}+\alpha_{7}site_{i}+\alpha_{8}race_{i}+ \varepsilon_{i}, CBCL_{Externalizing}\sim Gamma$$

This model explained 36.9% of the variance, with a GCV of 0.031.

The GCV validation indicated successful convergence after 1000 iterations (k-index > 1, *p* > 0.05), confirming the model's robustness.

**Table S3. The model selections in the relationships with the broad externalizing problems in GAMs.**

| **GAMs** | **Term: Imbalance score** | **Term: Age** | **Term: Sex (male)** | **Term: Imbalance score * sex** | **Term: Site** | **Model Comparison** |
| --- | --- | --- | --- | --- | --- | --- |
| Reduced model | **B = 1.800, SE = 0.768,**  ***p* = 0.019*** | **B = -0.526, SE = 0.137,**  ***p* < 0.001***** | **B = 0.716, SE = 0.173,**  ***p* < 0.001***** | - | *edf* = 0.003, *F* = 0.003,  *p* = 0.316 | - |
| Model 2 | ***edf* = 2.851, *F* = 2.909,**  ***p* = 0.025*** | **B = -0.527, SE = 0.137,**  ***p* < 0.001***** | **B = 0.724, SE = 0.173,**  ***p* < 0.001***** | - | *edf* = 0.042, *F* = 0.044,  *p* = 0.309 | *F* (2.696, 9363) = 2.575,  *p =* 0.058 (.vs reduced model) |
| Model 3 | **B = 2.190, SE = 0.799,**  ***p* = 0.006**** | **B = -0.491, SE = 0.138,**  ***p* < 0.001***** | **B = 0.711, SE = 0.172,**  ***p* < 0.001***** | - | ***F* (21, 9339.7) = 3.367,**  ***p* < 0.001***** | ***F* (20.994, 9363) = 3.418, *p <* 0.001***** (.vs reduced model) |
| Model 4 | **B = 0.049, SE = 0.017,**  ***p* = 0.004**** | **B = -0.011, SE = 0.003,**  ***p* < 0.001***** | **B = 0.015, SE = 0.004,**  ***p* < 0.001***** | - | ***F* (21, 9339.7) = 3.367,**  ***p* < 0.001***** | **GCV = 0.0311** (.vs model 3) |
| Model 5 | **B = 0.049, SE = 0.017,**  ***p* = 0.004**** | ***edf =* 1.003*, F* = 13.2,**  ***p* < 0.001***** | **B = 0.015, SE = 0.004,**  ***p* < 0.001***** | - | ***F* (21, 9339.7) = 3.367,**  ***p* < 0.001***** | ***F* (0.006, 9342) = 0.289,**  ***p =* 0.018*** (.vs model 4) |
| Model 6 | *-* | ***edf =* 1.002*, F* = 12.847,**  ***p* < 0.001***** | **B = 0.015, SE = 0.004,**  ***p* < 0.001***** | Male*: edf =*1.001*, F* = 1.061,  *p* = 0.303  Female: *edf =* 2.106*, F* = 1.061,  *p* = 0.003** | ***F* (21, 9339.7) = 3.367,**  ***p* < 0.001***** | *F* (2.723, 9342) = 2.680,  *p =* 0.051 |

### Supplement 4. Generalized Additive Models for Location, Scale and Shape: Trajectory of Dual-systems Imbalance

We applied generalized additive models for location, scale and shape (GAMLSS) to model the trajectory of the brain dual-system imbalance, allowing for flexible outcome distributions to accommodate non-Gaussian data and incorporate models for multiple moments of the outcome's distribution, including location (L), scale (S), and shape (S) parameters within the GAMLSS framework.

#### 5.1 Model distribution

Given the lack of prior knowledge to estimate the distribution of the imbalance score and the uncertainty about whether neuroimaging-based trajectories would align with known distributions, we evaluated all potential distribution families. Following standard practice, we fitted multiple distributions consistent with the outcome's characteristics and compared them using the Bayesian Information Criterion (BIC) (13). Given that the imbalance score ranged from -2 to 2, we evaluated distribution families with 1-4 parameters, all of which converged during GAMLSS fitting. The BIC analysis revealed that the Johnson SU distribution, with 4 parameters, provided the best fit (14).

#### 5.2 Centile normalization

The GAMLSS framework enables complex outcome distributions by parameterizing a distribution into multiple components. Each component is modeled as a regression with appropriate link functions (e.g., exponential links for sigma to ensure non-negative values for variance). We modeled the imbalance score as a function of four parameters (μ, σ, ν, τ), with each component potentially including distinct covariates.

Model specifications indicated the inclusion of a site-level random effect. Specifically, within the regression components, a random intercept was included per study, with uncorrelated random effects across components. The model structure was as follows:

$$Imbalance Score \sim JSU(\mu,\sigma,\upsilon,\tau)$$

$$g_{\mu}\left( \mu\right)=X_{\mu}\beta_{\mu}+{Z_{\mu}\gamma}_{\mu}+\sum_{i} s_{\mu,i}(x_{i})$$

$$g_{\sigma}\left( \sigma\right)=X_{\sigma}\beta_{\sigma}+{Z_{\sigma}\gamma}_{\sigma}+\sum_{i} s_{\sigma,i}\left( x_{i} \right)$$

$$g_{\upsilon}(\nu)=X_{\upsilon}\beta_{\upsilon}+{Z_{\upsilon}\gamma}_{\upsilon}+\sum_{i} s_{\upsilon,i}(x_{i})$$

$$g_{\tau}(\tau)=X_{\tau}\beta_{\tau}+{Z_{\tau}\gamma}_{\tau}+\sum_{i} s_{\tau,i}(x_{i})$$

In these, each component equation included: typical covariates and coefficients, 𝑋 and 𝛽; random-effects, 𝛾 (which may include covariates, 𝑍); and non-parametric smoothing functions (s) applied to covariates.

#### 5.3 Longitudinal centiles

Using the cumulative distribution function (CDF), we calculated longitudinal centiles for individuals with multiple observations. We hypothesized that within-subject covariance would be smaller than between-subject covariance, implying stability in an individual’s centile over time. We used the interquartile range (IQR) to summarize the stability of centiles across observations, making this metric comparable for participants with varying numbers of longitudinal data points. Given the variable number of longitudinal data-points available for different participants, we chose to use a measure that was consistent for participants who only had 2 observations as well as for participants with more than 2 observations. The median of IQR was 0.49 < 0.5, showing a stable trend (11).

#### 5.4 Model evaluation

The final model was as follows:

$$Imbalance Score \sim JSU(\mu,\sigma,\upsilon,\tau)$$

$$\mu=\alpha_{\mu}+\alpha_{\mu,sex}+\alpha_{\mu,race}+\alpha_{\mu,ethnic}+\beta_{\mu,1}{age}^{-2}+\gamma_{\mu,site}$$

$$log\left( \sigma\right)=\alpha_{\sigma}+\alpha_{\sigma,sex}+\alpha_{\sigma,race}+\beta_{\mu,1}age+\gamma_{\sigma,site}$$

$$\nu=\alpha_{\nu}$$

$$log(\tau)=\alpha_{\tau}$$

We evaluated model fit using traditional QQ-plots and residual distribution measures. The residuals followed a normal distribution (15), with a skewness of 0.0041, kurtosis of 3.016, and a Filliben correlation coefficient of 0.999 (FIGURE S1.B).

The worm plot showed that confidence intervals crossed the zero line, indicating normally distributed residuals (FIGURE S1.A) (16). The worm plot, which is a collection of Detrended QQ-plots across age groups, suggested useful modifications to the model based on the data points’ worm-like pattern.

#### 5.5 Model sensitivity analyses

To assess the robustness and reliability of the optimized GAMLSS models, we conducted several sensitivity analyses, including leave-one-study-out (LOSO) and 10-fold cross-validation.

For 10-fold cross-validation, we split the sample into a training set (70%) and a validation set (30%). The GAMLSS *gamlssCV()* function was used to fit the model and estimate the global deviance for the validation set (17), resulting in a global deviance of -318.8648, indicating excellent fit.

We further tested the model's reliability by performing a LOSO analysis, where each participant was iteratively excluded, model parameters were re-estimated, and fitted trajectories were extracted. This analysis confirmed that the model's reliability was not skewed by any individual participant.

### Supplement 5. Sensitivity Analyses

We conducted five sensitivity analyses to assess the robustness of our findings. First, we examined the trajectory of dual-systems imbalance and associations between imbalance scores and psychopathology scores using alternate dual-systems imbalance scores definitions based on the average deviation score (ADS) for surface area (SA), surface thickness (ST) and subcortical volume (SV) derived from a normative modeling framework. Second, we identified outliers in brain morphology features, which contributed to extreme imbalance scores potentially due to inaccurate segmentation or neurodevelopmental disorders. Outliers were removed to minimize bias, and all analyses were repeated.

#### 6.1 Outlier Removal

For each psychological measure, participants whose performance deviated more than three standard deviations (SD) from mean of the whole sample were considered as outliers and thus were excluded from further analysis, yielding a final sample of N = 22,019.

To determine the optimal neural correlate for the cognitive-control system, we assessed the predictive power of three surface morphology features using changes in marginal scores (${\Delta R}_{M}^{2}$) within LMMs. The SA (β = -0.093, SE = 0.01, *p* < 0.001, ${\Delta R}_{M}^{2}=7.363\times{10}^{-3}$) and GMV (β = -0.077, SE = 0.01, *p* < 0.001, ${\Delta R}_{M}^{2}$= 5.397×10^-3^) were both significantly associated with cognitive control, whereas ST (β = 0.013, SE = 0.009, *p* = 0.15) not significant, as illustrated in Figure S3. The SA exhibited the strongest predictive power for cognitive control, aligning with our main finding. Consequently, DLPFC SA was selected as the primary correlate of cognitive control for subsequent analyses.

As in the main analysis, an imbalance score was calculated as the difference of VS SV minus DLPFC SA then divided by their mean, with values ranging from -2 to 2. Structure-function analyses confirmed the validity of brain morphology features. DLPFC SA significantly predicted cognitive control, and VS GMV was significantly associated with sensation-seeking (β = 0.02, SE = 0.008, *p* < 0.05), as illustrated in Figure S2 (ABCD). Imbalance scores demonstrated high reliability (ICC = 0.935, 95% CI: 0.931 to 0.94).

The trajectory analysis revealed that cognitive control matures faster than the socioemotional system during early adolescence, with a balance emerging around age 13. Positive imbalance scores reflected the prematurity of the socioemotional system. The variance of imbalance scores increased during adolescence as well as growth rate, consistent with the main analysis.

When assessing psychopathology, we found similar patterns to the main analysis. The relationship with broad internalizing showed a significantly non-linear pattern ($F=2.34, p< 0.05$), with an initial decrease followed by an increase, explaining 35% of the variance. The relationship with broad externalizing showed a significantly positive association (B = 0.02, SE = 0.0068, *p* < 0.01), explaining 36.6% of the variance, aligning with the main analyses. Non-linear age effects were significant for both internalizing $(age:F=5.353, p<0.05) and externalizing$ $(age:F=7.362, p<0.05)$ dimensions.

#### 6.2 Average deviation score (ADS)

Sex-specific normative models for regional ST, SA, and SV were developed using a large multisite sample of 37,407 healthy individuals, made available by the ENIGMA Lifespan Group through Centile Brain, an open-access web portal (18). The normative model details were in open science ([CentileBrain](https://centilebrain.org/" \l "/)).

We applied these models to estimate the normative deviation for each measure in our sample. Z-scores were computed for each region, indicating how much a participant's morphometric measures deviated from the population mean. The ADS values were not weighted by regional size to enhance reproducibility. Positive or negative z-scores indicated higher or lower values compared to the normative mean, respectively (19).

The ADS for ST (β = 0.0406, SE = 0.009, *p* < 0.001) was significantly associated with cognitive control, while the ADS for SA (β = -0.000604, SE = 0.0091, *p* = 0.947) was not. Consequently, imbalance scores were calculated as the difference between the ventral striatum (VS) SV ADS and dorsolateral prefrontal cortex (DLPFC) ST ADS, with values ranging from -4 to 4. Positive values indicated greater cognitive control system maturity relative to the socioemotional system. The imbalance scores demonstrated high reliability (ICC = 0.926, 95% CI: 0.921 to 0.931).

$${Imbalance Score}_{ADS}={ADS(VS)}_{Volumn, scale}-{ADS(DLPFC)}_{Thickness, scale}$$

Structure-function analyses validated the validity of morphological features. DLPFC ST ADS was a significant statistical predictor of cognitive control, and VS SV was significantly associated with sensation-seeking (β = 0.022, SE = 0.01, *p* < 0.05), as illustrated in Figure S3.

The GAMLSS model indicated that the ADS imbalance score was negative and decreased over time, suggesting that cognitive-control system maturation outpaced socioemotional system during ages from 9 to 16. The variance of ADS of imbalance scores gradually decreased during adolescence, and its growth rate gradually decreased. These findings may have differed from the main analysis due to: 1) different neural correlates of cognitive-control system; 2) the non-weighted nature of the ADS calculation; 3) the use of ADS in a trajectory context rather than for case-control comparisons, as it was originally intended. Future research should consider the appropriate use of the ADS in trajectory modeling.

Therefore, when evaluating psychopathology, we found that the imbalance score was not significantly associated with internalizing problems but showed a significant negative association with broad externalizing behaviors (B = -0.0067, SE = 0.002, *p* < 0.001), consistent with the Dual Systems model, which links greater imbalance to more risky behaviors. This consistency in externalizing outcomes, despite some discrepancies, underscores the robustness of the imbalance score in capturing dual-system dynamics.

#### 6.3 Human Connectome Project in Development (HCP-D)

We applied the independent sample of HCP-D to validate the generalizability and robustness of our results. Participants lacking brain T1 image data (N = 27), psychopathological measurements (N = 22) or demographic information (N =12) were excluded from the analysis, resulting in a final sample size of N = 591.

**GAMLSS: trajectory of imbalance score**

The best distribution for the HCP-D sample was identified as the logistic distribution ("LO" in the GAMLSS package), characterized by two parameters. Following the workflow outlined in Supplement 3, we derived the trajectory, variance, and growth rate of the imbalance score, which aligned with our findings from the ABCD study.

**Associations with Psychopathological Dimensions**

The models mirrored our main analysis in the ABCD sample, incorporating sex, age, race, ethnicity, familial income, parental highest education, and marital status as fixed effects, alongside other psychopathological dimensions. Given that each family in the HCP sample contained only one child, we excluded family effects from our models and included site as a random effect. The model indicated boundary singular fit, suggesting insufficient variance in the random effect to explain random variability. Upon reviewing site information—comprising four sites with similar child numbers—we posited that this contributed to poor model fitting. Consequently, we removed the random effect to refine the analysis.

Regarding the relationship between the imbalance and broad internalizing scores, the GAM revealed that the smooth term for the imbalance score was not significant. However, this may not contradict our main analysis, as the smooth term was substantially influenced by sample size. Supporting this, polynomial models demonstrated a significant quadratic effect of the imbalance score (*p* < 0.05), indicating a non-linear relationship.

For broad externalizing scores, we observed similar results. We modeled four GAMs to examine associations between the imbalance and broad externalizing scores, as well as the effects of age and the interactions between the imbalance scores and sex. Model1 included all linear covariates (including broad internalizing) and treated the imbalance score as a linear effect, yielding a good fit (GCV = 53.071, R^2^(adj) = 0.351) with a significant association between the imbalance and externalizing scores (B = 1.886, SE = 0.726, *p* < 0.05). Model 2 incorporated a smooth term for the imbalance score, which proved significant ($F=3.367, p<0.05$), while maintaining a comparable fit (GCV = 52.704, R^2^(adj) = 0.353). This suggests that the linear effect was sufficient, and the inclusion of a smooth term might have introduced overfitting. Models 3 explored the smooth effects of age, while model 4 examined the interaction between the imbalance scores and sex. However, model comparisons did not indicate significant improvements ($Model3: GCV=52.52, R^{2}(adj)=0.355, \chi^{2}\left( 1.47 \right)=2.705, p=0.083;Model4: GCV=53.535, R^{2}(adj)=0.342$). Thus, the model demonstrated a significant linear association between the imbalance scores and externalizing behaviors.

Interestingly, through the U-shape relationship between the imbalance and broad internalizing scores, it is likely necessary to consider additional brain systems not included in the Dual Systems model to fully account for development from preadolescence to adulthood. Expanding the framework to include additional systems, such as the triadic model (20,21) could interpret the non-linear pattern with internalizing and offer a more complete understanding of adolescent development. For example, the role of the amygdala in linking cognitive-control and reward systems warrants further investigation (22), as it may provide additional insights into psychopathology emerging during adolescence (23).

#### 6.4 Global Imbalance

The observed transdiagnostic effects of the imbalance score and its alignment with the Dual Systems Model raise the critical question of whether these effects are driven by global anatomical measures or specific regional contributions from the VS and DLPFC. To address this, we constructed a global imbalance score using total surface area (corresponding to the SA of DLPFC) and global subcortical volume (corresponding to the SV of VS):

$$Global Imbalance Score={Subcortical Volumn}_{scale}-{Surface Area}_{scale}$$

**Developmental Trajectory of the Global Imbalance**

The best distribution for the global imbalance score was identified as the Skew Student t distribution ("SST" in the *GAMLSS* package). This model is characterized by four parameters: the mean (*μ*), standard deviation (*σ*), skewness (ν), and kurtosis (τ) (24). Following the workflow outlined in Supplement 3, we derived the trajectory and growth rate of the global imbalance score. The final model specification was:

$$Global Imbalance Score \sim SST(\mu,\sigma,\upsilon,\tau)$$

$$\mu=\alpha_{\mu}+\alpha_{\mu,sex}+\alpha_{\mu,race}+\beta_{\mu,1}age+\gamma_{\mu,site}$$

$$log\left( \sigma\right)=\alpha_{\sigma}+\alpha_{\sigma,sex}+\alpha_{\sigma,ethni}+\beta_{\mu,1}age+\gamma_{\sigma,site}$$

$$log(\nu)=\alpha_{\nu}$$

$$log(\tau)=\alpha_{\tau}$$

The model evaluations are shown in Figure S4A-B, and the results demonstrated good model fit. The global imbalance trajectory revealed a linear increase in scores with age, maintaining positive values between 9 and 16 years (Figure S4C). This contrasts with the developmental trajectory of the primary imbalance score, which showed decreasing values. Additionally, the variance of the global imbalance score increased during this period.

**Associations with Psychopathological Dimensions**

The models mirrored our main analysis, incorporating sex, age, race, ethnicity, familial income, parental highest education, and marital status as fixed effects, alongside other psychopathological dimensions. All GAMs were analyzed in the *mgcv* package in R 4.4.1.

Initial LMM analyses indicated a significant positive relationship between the global imbalance score and externalizing symptoms (B = 6.999, SE = 1.268, *p* < 0.001) and explained 48.9% of the variance. To explore potential non-linearities, GAMs were applied. The Model 1 showed that the global imbalance score was linearly significant (B = 8.608, SE = 1.505, *p* < 0.001, $GCV=66.946, R^{2}\left( \mathrm{adj} \right)=0.374$). The Model 2 showed a significant non-linear pattern of the global imbalance score ($GCV=66.945, R^{2}\left( \mathrm{adj} \right)=0.374, edf=1.702, F=15.03, p<0.001$). The model comparison showed no significant difference ($F\left( 1.161, 9342 \right)=1.337, p=0.253$). Given the rate of explanation for variation, we finally adopted the LMM to portray their relationships.

For internalizing symptoms, LMM and segmented polynomial models found no significant associations (LMM: B = -0.979, SE = 1.330, *p* = 0.462, R^2^ = 48.8%; quadratic: B = 11.43, SE = 0.872, *p* = 0.197; cubic: B = -8.740, SE = 8.719, *p* = 0.316). Model 1 of the GAM showed the global imbalance score was not significantly associated with the internalizing scores (B = -0.828, SE = 1.572, *p* = 0.598, $GCV=72.753, R^{2}\left( \mathrm{adj} \right)=0.365$), and Model 2 showed the non-linear pattern of the global imbalance score was not significant ($GCV=72.736, R^{2}\left( \mathrm{adj} \right)=0.365, edf=2.799, F=1.082, p=0.309$), while the model comparison showed no significant difference ($F(2.572, 9342)=2.273, p=0.088$). The poor overall model fit and lack of significant differences between linear and non-linear models indicated that global imbalance is a poor predictor of internalizing symptoms.

The global imbalance scores significantly predicted externalizing symptoms, suggesting that global cortical and subcortical morphological differences may contribute to these behaviors. However, the poor predictive value for internalizing symptoms underscores the importance of regional specificity in the primary imbalance score. These results suggest that the observed transdiagnostic effects are driven by the specific neural systems we chose for the imbalance score rather than global anatomical features.

#### 6.5 Alternative Imbalance Scores

Emerging evidence suggests that neural systems connecting the prefrontal cortex (PFC) to mesolimbic regions, including the amygdala and striatum, mediate reward processing and sensation-seeking behaviors (25). The amygdala, a key structure in the impulsive system, has been linked to risk-taking and externalizing problems (26,27), with reduced amygdala gray matter often reported in addiction (28,29).

To assess the specificity of our imbalance score, we calculated an alternative imbalance score using the difference between the amygdala and DLPFC. This allowed us to evaluate whether observed effects extend to other regions within the socioemotional and cognitive-control systems.

$$Alternative Imbalance Score={Amygdala volume}_{scale}-{DLPFC area}_{scale}$$

**Developmental Trajectory of the Alternative Imbalance**

The best distribution for the alternative imbalance score was identified as the Skew Student t distribution ("SST" in the *GAMLSS* package). This model is characterized by four parameters: the mean (*μ*), standard deviation (*σ*), skewness (ν), and kurtosis (τ) (24). Following the workflow outlined in Supplement 3, we derived the trajectory and growth rate of the alternative imbalance score. The final model specification was:

$$Alternative Imbalance Score \sim SST(\mu,\sigma,\upsilon,\tau)$$

$$\mu=\alpha_{\mu}+\alpha_{\mu,sex}+\alpha_{\mu,race}+\beta_{\mu,1}age+\gamma_{\mu,site}$$

$$log\left( \sigma\right)=\alpha_{\sigma}+\alpha_{\sigma,sex}+\beta_{\mu,1}age+\gamma_{\sigma,site}$$

$$log(\nu)=\alpha_{\nu}$$

$$log(\tau)=\alpha_{\tau}$$

The model evaluations are shown in Figure S5A-B, and the results demonstrated good model fit. The global imbalance trajectory revealed a non-linear increase in scores with age, maintaining negative values between 9 and 16 years (Figure S5C).

**Associations With Sensation-Seeking**

A linear mixed model (LMM) was conducted to evaluate the relationship between sensation-seeking and SV of the amygdala. The reduced model included socio-demographic covariates, with site and family nested within site as random effects, while the full model added amygdala SV as a fixed factor. Results showed a significant association between amygdala SV and sensation-seeking (β = 0.641, SE = 0.284, 95%CI: 0.084 to 1.198, *p* = 0.024, ${\Delta R}_{M}^{2}$= 5.926×10^-4^).

**Associations With Externalizing Symptoms**

The alternative imbalance score was positively associated with broad externalizing symptoms in LMMs (B = 2.898, SE = 0.752, *p* < 0.001). Model diagnostics indicated potential non-linearity in residuals, which was further supported by segmented polynomial models (See Table S6).

Table S6. The relationships between the broad externalizing and the imbalance scores in segmented polynomial models.

| **Segmented Polynomial models** | **Imbalance score, 1** | **Imbalance score, 2** | **Imbalance score, 3** | **Imbalance score, 4** | **Imbalance score, 5** |
| --- | --- | --- | --- | --- | --- |
| 2 | **B = 35.08, SE = 8.981, *p* < 0.001***** | B = -12.01, SE = 8.313, *p* = 0.148 | - | - | - |
| 3 | **B = 34.91, SE = 8.980, *p* < 0.001***** | B = -12.03, SE = 8.312, *p* = 0.148 | B = 11.53, SE = 8.206, *p* = 0.160 | - | - |
| 4 | **B = 34.94 SE = 8.980, *p* < 0.001***** | B = -12.08, SE = 8.313, *p* = 0.146 | B = 11.46, SE = 8.207, *p* = 0.162 | B = 6.678, SE = 8.182, *p* = 0.414 | - |
| 5 | **B = 34.94, SE = 8.981, *p* < 0.001***** | B = -12.08, SE = 8.313, *p* = 0.146 | B = 11.46, SE = 8.207, *p* = 0.162 | B = 6.682, SE = 8.183, *p* = 0.414 | B = -39.83, SE = 8.184, *p* = 0.961 |

Table S7. The relationships between the broad externalizing and the imbalance scores in GAMs.

| **GAMs** | **Term: Imbalance score** | **Term: Age** | **Term: Sex (male)** | **Term: Imbalance score * sex** | **Term: Site** | **Model Comparison** |
| --- | --- | --- | --- | --- | --- | --- |
| Reduced model | B = 2.934, SE = 0.758,  *p* < 0.001*** | B = -0.562, SE = 0.137,  *p* < 0.001*** | B = 0.740, SE = 0.172,  *p* < 0.001*** | - | *edf* = 0.003, *F* = 0.003,  *p* = 0.315 | - |
| Model 2 | B = 3.345, SE = 0.830,  *p* < 0.001*** | B = -0.522, SE = 0.138,  *p* < 0.001*** | B = 0.737, SE = 0.172,  *p* < 0.001*** | - |  | *F* (20.994, 9363) = 3.383,  *p <* 0.001*** (.vs reduced model) |
| Model 3 | *edf* = 1.761, *F* = 7.857,  *p* < 0.001*** | B = -0.521, SE = 0.138,  *p* < 0.001*** | B = 0.742, SE = 0.172,  *p* < 0.001*** | - |  | *F* (1.241, 9342) = 1.798, *p =* 0.178 (.vs model 2) |
| Model 4 | B = 0.070, SE = 0.018,  *p* < 0.001*** | B = -0.011, SE = 0.003,  *p* = 0.003** | B = 0.015, SE = 0.003,  *p* < 0.001*** | - |  | - |
| Model 5 | B = 0.070, SE = 0.018,  *p* < 0.001*** | *edf =* 1.002*, F* = 14.91,  *p* < 0.001*** | B = 0.015, SE = 0.003,  *p* < 0.001*** | - |  | *F* (0.005,9342) = 0.263,  *p =* 0.018* |
| Model 6 | - | *edf =* 1.003*, F* = 13.996,  *p* < 0.001*** | - | Male*: edf =* 1.448*, F* = 3.115,  *p* = 0.034*  Female: *edf =* 1.004*, F* = 5.666,  *p* = 0.017* |  | - |

The model selections in GAMs are in Table S7 and model5 is the final pattern, in which the alternative imbalance scores positively associated with the broad externalizing problems (B = 0.070, SE = 0.018, *p* < 0.001) and could explain 37.8% of the variance.

**Associations with Internalizing Symptoms**

For internalizing symptoms, the results were less consistent. In the LMM, the alternative imbalance score was not significantly associated with broad internalizing problems (B = -1.101, SE = 0.784, *p* = 0.161). However, the segmented polynomial models did reveal some evidence supporting this relationship (See Table S8), indicating that quadratic terms may capture non-linear effects.

Table S8. The relationships between the broad internalizing and the alternative imbalance scores in segmented polynomial models.

| **Segmented Polynomial models** | **Imbalance score, 1** | **Imbalance score, 2** | **Imbalance score, 3** | **Imbalance score, 4** | **Imbalance score, 5** |
| --- | --- | --- | --- | --- | --- |
| 2 | B = -14.79, SE = 9.060, *p* = 0.102 | B = 23.79, SE = 8.574, *p* = 0.005** | - | - | - |
| 3 | B = -14.88, SE = 9.060, *p* = 0.101 | B = 23.80, SE = 8.574, *p* = 0.005** | B = 8.281, SE = 8.493, *p* = 0.329 | - | - |
| 4 | - | - | - | - | - |
| 5 | - | - | - | - | - |

Interestingly, GAMs did not find significant smoothing effects (Table S9) for the alternative imbalance score in relation to internalizing symptoms, contrasting with the quadratic significance observed in polynomial models. This seeming discrepancy may arise from differences in how the models handle random effects. Specifically, the LMM and polynomial models accounted for site and family as random effects, while the GAM reduced familial effects by randomly selecting one participant per family. The lack of significant results in the GAM suggests that the relationship between the alternative imbalance score and internalizing problems is less robust and potentially influenced by methodological choices.

Table S9. The relationships between the broad externalizing and the alternative imbalance scores in GAMs.

| **GAMs** | **Term: Imbalance score** | **Term: Age** | **Term: Sex (male)** | **Term: Imbalance score * sex** | **Term: Site** | **Model Comparison** |
| --- | --- | --- | --- | --- | --- | --- |
| Reduced model | B = -1.272, SE = 0.791,  *p* = 0.107 | B = 0.383, SE = 0.143,  *p* = 0.007** | B = 0.843, SE = 0.179,  *p* < 0.001*** | - | *edf* = 0.740, *F* = 2.856,  *p* = 0.049* | - |
| Model 2 | *edf* = 2.337, *F* = 2.639,  *p* = 0.0459* | B = 0.380, SE = 0.143,  *p* = 0.008** | B = 0.834, SE = 0.179,  *p* < 0.001*** | - | *edf* = 0.743, *F* = 2.897,  *p* = 0.048* | *F* (1.990, 9362.1) = 3.739,  *p =* 0.024* (.vs reduced model) |
| Model 3 | *edf* =2.342, *F* = 2.198,  *p* = 0.084 | B = 0.455, SE = 0.144,  *p* = 0.002** | B = 0.852, SE = 0.179,  *p* < 0.001*** | - |  | *F* (20.075, 9360) = 4.711, *p <* 0.001*** (.vs model 2) |

**Comparative Interpretation of the Imbalance and Alternative Imbalance Scores**

We observed that the alternative imbalance score exhibited a consistent and significant relationship with externalizing symptoms, while its association with internalizing problems was weaker and less stable. Additionally, the explanatory power and model fit of the alternative imbalance score for externalizing symptoms was lower compared to the original imbalance score (F(0.371, 9339.9) = 14.523, *p* = 0.005), which specifically focused on the VS-DLPFC axis.

This pattern underscores the unique contribution of the DLPFC and VS to the transdiagnostic associations between dual-systems imbalance and psychopathology. The original imbalance score captures specific neurodevelopmental mechanisms tied to these regions, which appear to have stronger and more stable transdiagnostic associations than those derived from the amygdala-DLPFC axis. These findings suggest that the predictive utility of imbalance scores is enhanced when anchored to well-defined and theory-driven neuroanatomical substrates.

### Supplementary 6 The Rationale to choose the dorsolateral prefrontal cortex and ventral striatum

The selection of the DLPFC and VS aligns well with dual systems models, which delineate distinct neural substrates for cognitive control and socioemotional processing. These regions correspond to the cognitive control and socioemotional system, respectively. Although theories suggest that dual systems encompass additional brain regions, the DLPFC and VS were prioritized as central and representative regions, serving as effective proxies for their respective systems. Below, we elaborate on the significance and unique contributions of these brain regions.

#### Socioemotional System: Ventral Striatum

Within Dual Systems Models, the socioemotional system has been proposed to be primarily localized in the striatum, as well as the medial and orbitofrontal cortices(30–32). However, developmental neuroscience research has predominantly concentrated on developmental differences within the striatum, particularly the ventral striatum (nucleus accumbens), which is proposed as a central structure in the reward system (23,33,34) Both functional and structural evidence robustly implicate the VS in reward processing and reactivity (35,36)

The striatum comprises the caudate nucleus, putamen, and nucleus accumbens, each of which contributes uniquely to socioemotional processing through corticostriatal connections (37). MRI studies have revealed that striatal morphology is altered in neurodegenerative conditions, providing critical insights into the links between genotype, structure, and function. Specifically, the morphology of the VS, including volumetric measures, has been linked to reward sensitivity and motivational processes (33,38). Notably, the ventral striatum exhibits the greatest developmental differences and contributes most significantly to multivariate age predictors (39). Structural analyses have associated VS volume with individual differences in sensation-seeking and reward-related behaviors, further validating its role as a marker for the socioemotional system (39). Advanced MRI methodologies, including T2* imaging, have demonstrated that the morphology of the striatum is linked to neurophysiological maturation. For instance, T2* signal variations in the VS indicate changes in tissue–iron concentration, which are associated with dopamine function and myelination (40). These neurophysiological changes support the maturation and proliferation of the dopamine system and the myelination of cortico-striatal connections observed in animal models of adolescent development (30,39). Such findings highlight the VS's potential as a biomarker for structural and functional assessments related to socioemotional reactivity and reward processing.

Moreover, the voxel-wise distribution of feature weights from multivariate support vector regression indicates that neurophysiological maturation of the striatum is most strongly influenced by the continued maturation of the VS, including the nucleus accumbens and ventromedial portions of the caudate and putamen, into adulthood. During adolescence, the VS exhibits peak functional reactivity to reward stimuli under certain incentive contexts and is associated with risk-taking behavior (40,41). This region is highly dopamine-innervated and is a central component of the fronto-striatal dopamine reward pathways, hypothesized to underlie sensation-seeking and risk-taking behavior. Increases in tissue–iron concentration in the VS may be mechanistically related to adolescent behavior and striatal reward reactivity through associations with dopamine receptor expression, transporter function, excitability, and myelination within cortico-ventral striatal pathways (42).

In summary, the selection of the VS as a structural measure is justified by its integral role within corticostriatal circuits, its unique developmental trajectory, and its consistent association with reward-related behaviors and socioemotional reactivity. By focusing on VS volume, our study leverages a structural measure that is both theoretically and empirically validated within the context of dual systems models.

#### Cognitive Control System: Dorsolateral Prefrontal Cortex

The prefrontal cortex undergoes significant progressive and regressive changes throughout development, with synaptic pruning in the DLPFC continuing into the second decade of life (32). The DLPFC plays a pivotal role in executive functions, including working memory, cognitive flexibility, planning, and inhibitory control (1,43). These executive functions are essential for the deliberative processes that underpin cognitive control, enabling individuals to regulate impulses and make informed decisions. Neuroimaging studies consistently demonstrate DLPFC activation during tasks requiring cognitive control, such as the Stroop task, Go/No-Go tasks, and anti-saccade tasks. Importantly, the DLPFC serves as the brain's entry node for processing information relevant to cognitive and executive functions (44,45).

Developmental differences in DLPFC activation have been extensively investigated using approaches like performance on the oculomotor delayed response (ODR) task, which manipulates delays to assess working memory across age groups. In a functional MRI study examining children (10–13 years), adolescents (14–17 years), and adults (18–30 years), the DLPFC was recruited across all groups, but the magnitude of right DLPFC activation followed an inverted U-shaped trajectory, peaking during adolescence (46). These results likely reflect developmental immaturities in processes underlying cognitive control. Poorer performance by children was associated with greater reliance on basal ganglia and insula, whereas adults demonstrated a more distributed recruitment of temporal regions. Adolescents performed similarly to adults but displayed increased DLPFC recruitment, possibly reflecting greater effort. Studies of verbal working memory also show that adults recruit multiple frontal and parietal regions, with recruitment increasing systematically with cognitive load (47,48).

Similarly, an event-related design using the ODR task with varying delay periods (short: 2.5 seconds; long: 10 seconds) revealed that children and adolescents relied more on extended DLPFC circuitry during longer delays, while adults exhibited increased activation in specialized regions, including the inferior frontal gyrus (IFG) and parietal regions (49). During longer delay periods, children and adolescents relied on an extended circuitry including DLPFC while adults showed increased activation of parietal regions and IFG. These results indicate that again, younger individuals relied more on the DLPFC, while adults utilized seemingly more specialized regions that support better accuracy.

Studies where performance is equated have found that DLPFC activation, a region which supports working memory, response planning, and regulation needed for cognitive control, decreases with age (50). These results could also reflect immaturities in PFC structure, which could result in a more prolonged or extended computational process, generating increased activity compared to the adult system (51). These findings suggest that younger individuals may depend more heavily on the DLPFC to compensate for less efficient processing. Structural studies further link DLPFC volume and surface area with individual differences in cognitive control, underscoring its role as a critical marker of executive function.

The DLPFC and VS have emerged as central nodes within the dual systems framework, representing cognitive control and socioemotional systems, respectively. These regions have been identified as key players in numerous studies across developmental, structural, and functional domains. The integration of both structural and functional data provides a comprehensive understanding of these regions' contributions to the dual systems. The convergence of evidence across methodologies strengthens the validity of using these regions as proxies for their respective systems. While the DLPFC and the VS are selected as proxies, it is important to acknowledge that other regions could also contribute to these systems. To confirm the specificity and robustness of our selected markers, imbalance scores incorporating additional regions were constructed. The replication of findings in the HCP-D sample underscores the generalizability and robustness of using DLPFC and VS as neuroanatomical markers. We also constructed alternative imbalance scores using additional brain regions related to dual systems theories, such as the amygdala, which further support the appropriateness and specificity of the DLPFC and VS as markers.

### Supplementary7 The Rationale to choose brain structure rather than functional index

The personalized imbalance score we introduced utilized the VS volume and DLPFC surface area as proxies for the dual systems framework, with evidence of the neuroanatomy-behavior correlations supporting the validity of this selection. Theoretical justification for choosing the VS and the DLPFC as proxies for reward sensitivity and cognitive control, respectively, was detailed in Supplementary 6. Further, we investigated the criterion-related validity through the psychological/behavioral level measures, constructs such as sensation seeking and cognitive control were applied to, with significant results and medium effect sizes supporting the selection of these proxies.

Although functional measures, such as task-based or resting-state fMRI, are widely used in studies testing or extending dual systems theory, they are inherently state-dependent. However, the limitation was that the functional measures (e.g., fMRI activation) are state-dependent, influenced by temporary or contextual factors like mood, task engagement, or external stimuli. The morphometric features were biological stable, and the stable measures could serve as stable proxies for the developmental changes.

More importantly, anatomical measures are particularly effective for revealing individual differences and providing insights into behavior and psychopathology. The dynamic neurodevelopmental processes may include synaptic pruning, myelination, and cortical expansion, as well as the activity variance of the dopamine system (52), which may influence approach behaviors and preferences. For example, the immaturity of the DLPFC structure correlates with its reduced regulatory capacity during adolescence (53). These neurochemical underpinnings that are anchored in brain structural features offer biologically valid insights into individual differences, perhaps surpassing the transient nature of many functional states. The quantifiability of structural measures further enhances their utility. High-resolution neuroimaging techniques, such as MRI, provide precise and reliable measurements of brain anatomy across individuals and time points, allowing for the detection of subtle yet meaningful variations that underlie behavioral and psychological variability.

While considerable research has investigated psychological aspects of dual systems, trait-based individual differences may be driven by neurobiological substrates that interact with age-related developmental trajectories. Understanding better the underlying neurobiological mechanisms that drive states or abilities, rather than focusing solely on cognitive capacities or reward-seeking tendencies, is important (33). Neurochemical compounds, such as dopamine, may represent key drivers of these processes. However, many developmental studies are preclinical, with limited direct testing in humans, presenting ongoing challenges for translation and application. Thus, focusing on structural and neurobiological substrates to bridge the gap between preclinical findings and clinical applications may help provide a more robust framework for understanding adolescent neurodevelopment and its implications for psychopathology.

### Supplementary 8. The Rationale for Choosing Sensation Seeking and Cognitive Control

In the context of this study, we conceptualize "reward sensitivity" and "cognitive control" as core neurobiological constructs within the framework of Dual Systems Models. These constructs are operationalized through measurements of brain structure and function, while their psychological manifestations—"sensation seeking" and "cognitive control"—are assessed via subjective reports, capturing psychological states or tendencies. This delineation allows us to bridge neurobiological processes and observable behaviors within the context of developmental and psychopathological research.

**Sensation Seeking**

The Dual Systems Models employ the term "sensation seeking" as an umbrella label for interrelated constructs characterized by a propensity to "seek varied, novel, complex, and intense sensations and experiences and a willingness to take physical, social, legal, and financial risks for the sake of such experiences" (54). This construct has been consistently associated with heightened activity and recruitment of brain regions implicated in reward processing, particularly the VS. Evidence from human and animal studies supports this association, linking VS function to sensation-seeking behaviors across diverse contexts. Sensation seeking reflects the psychological expression of reward sensitivity, a hallmark of the socioemotional system, and provides a measurable proxy for the neurobiological underpinnings of this system.

**Cognitive Control**

In parallel, we use the term "cognitive control" to describe a cluster of related but distinguishable constructs that reflect an individual's ability to consciously regulate thoughts, emotions, or actions to achieve planned goals. Constructs such as impulse control, response inhibition, emotion regulation, and attentional control fall under this overarching label. Cognitive control has been robustly linked to the functioning of brain regions and systems subserving executive processes, most notably the lateral prefrontal cortex and lateral parietal regions (34). These regions collectively form the neurobiological foundation of the cognitive-control system, enabling top-down modulation of behaviors that counterbalance the socioemotional system's drive for immediate reward.

### FIGURE S1. Model Diagnosis

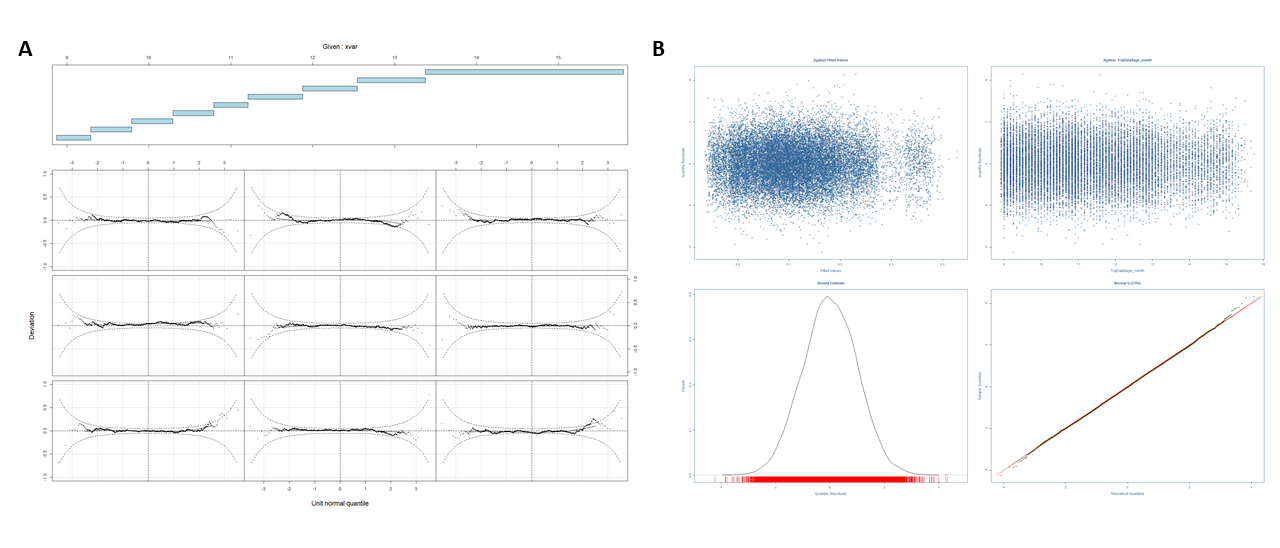
***Panel A*** depicts the worm plot from the GAMLSS model, illustrating that the confidence intervals cross the zero line, indicating residuals are approximately normally distributed. The worm plot presents detrended QQ plots across successive age groups, with the vertical axis reflecting the difference between observed and theoretical quantiles, forming the "worm-like" pattern characteristic of well-fitted models. ***Panel B*** presents the traditional QQ plot and residual analysis for the GAMLSS model fit. The residuals demonstrate adherence to a normal distribution, as evidenced by a skewness of 0.0041, kurtosis of 3.016, and a Filliben correlation coefficient of 0.999, confirming model fitting.

### FIGURE S2. Sensitivity Analysis of Outlier Removal and Independent Dataset of HCP-D

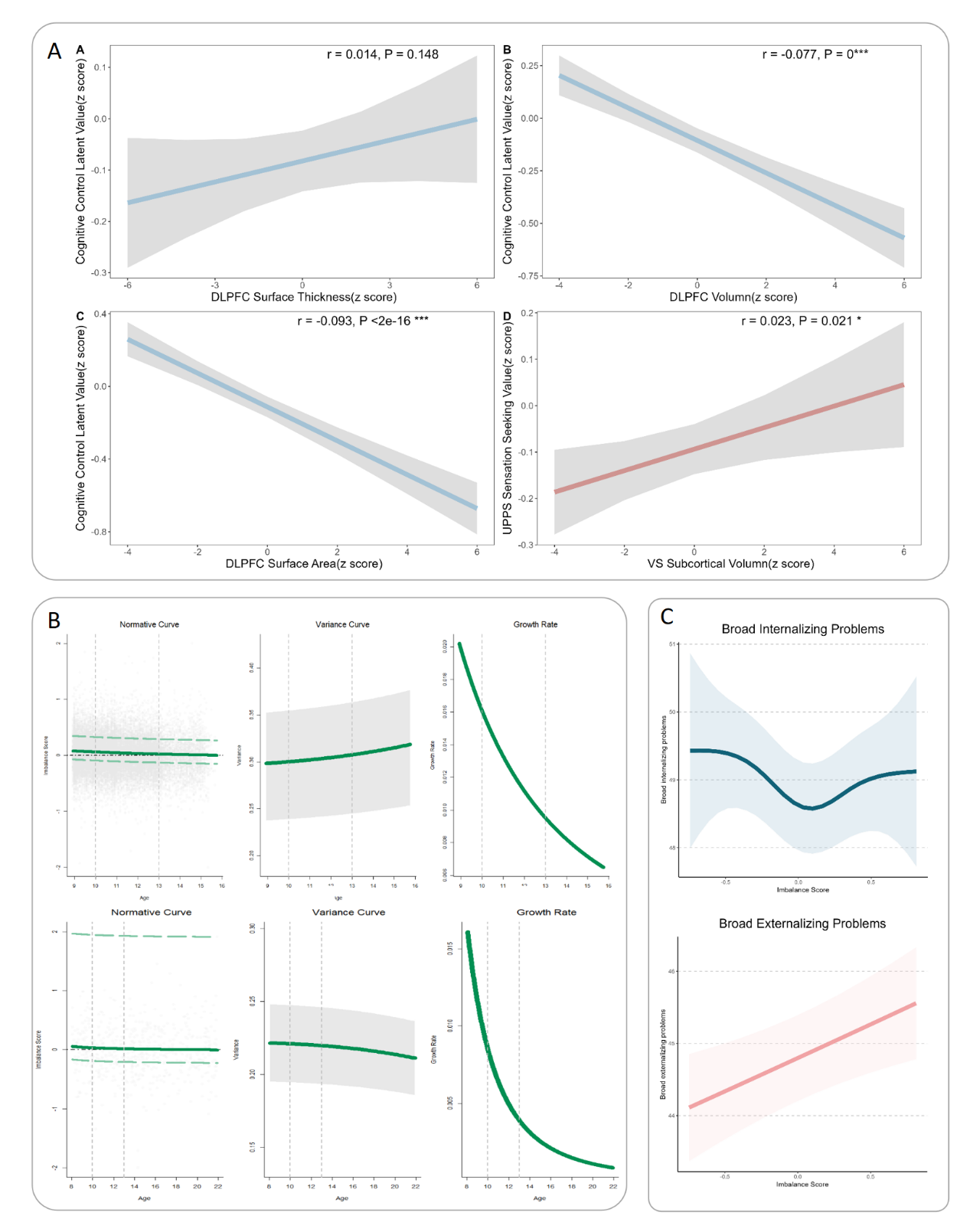

***Panel A*** illustrate the association between brain morphological features and psychological measures related to the dual-systems model after outlier exclusion. The surface area (SA) and volume of the dorsolateral prefrontal cortex (DLPFC) are significantly linked to cognitive control, while the subcortical volume (SV) of the ventral striatum (VS) is significantly related to sensation-seeking. Marginal R²calculations indicate that the SA of DLPFC is a primary neural correlate of the cognitive-control system. Although cognitive ability is highly polygenic and research to date suggests its cortical substrate is highly poly-regional, the positive relationship between the SA of DLPFC and cognitive control is consistent with published findings (55).

***Panel B*** displays the trajectory of imbalance scores, calculated as the difference between the SV of VS and the SA of DLPFC divided by their mean, with values ranging from -2 to 2. Scores range from -2 to 2, with higher values indicating greater maturity of the socioemotional system relative to the cognitive-control system. On the top are the results of outlier removal. The GAMLSS model fit for the imbalance trajectory, shown in the first column, reveals a positive score during early adolescence, which decreases over time, signifying early socio-emotional system maturity and gradual cognitive control development. Similar results are found in the HCP-D data. The second column shows the variance in the sample during adolescence, and the third column demonstrates the growth rate of the imbalance score.

***Panels C*** depicts the relationship between imbalance scores and psychopathological dimensions after outlier exclusion. The imbalance scores exhibit a significant nonlinear association with broad internalizing symptoms and a strong linear correlation with broad externalizing symptoms, in line with the primary analysis.

### FIGURE S3. Analysis using the Average Deviation Score

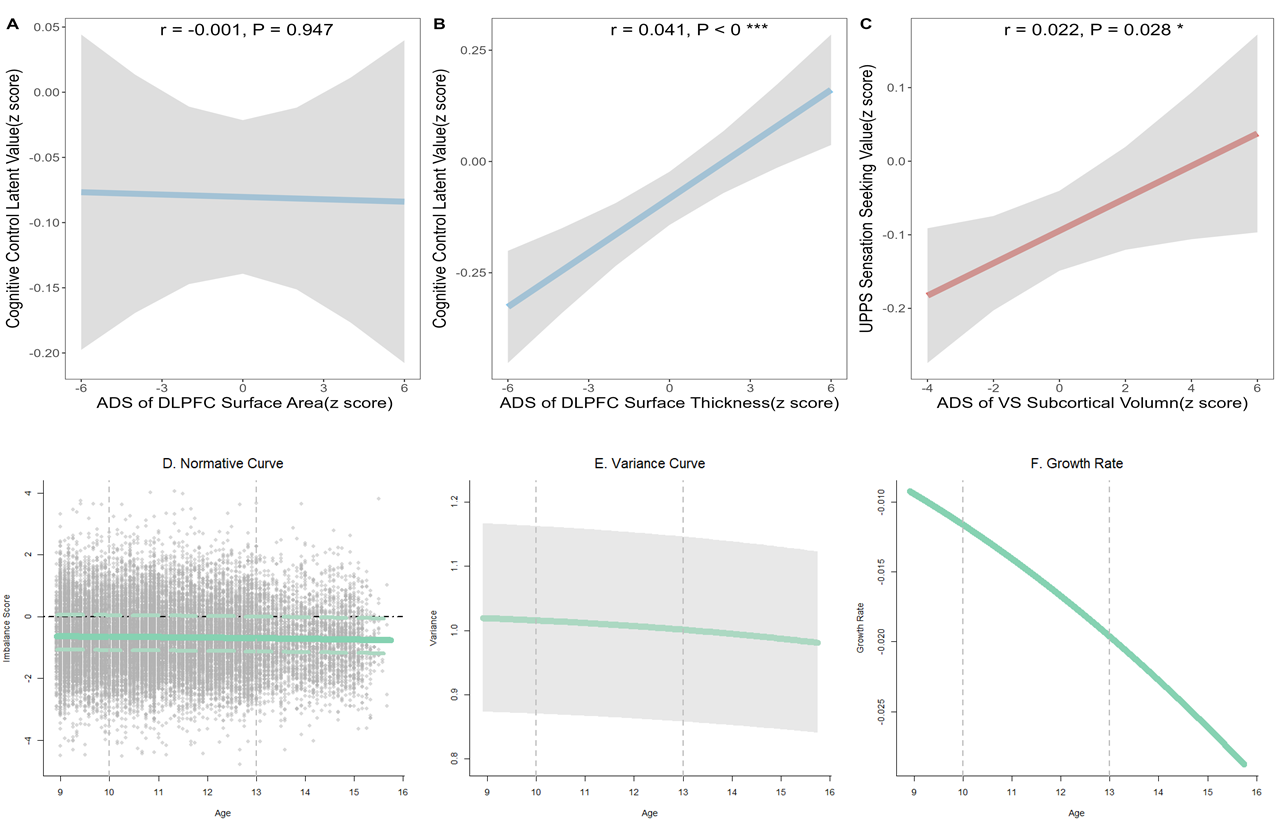

***Panels A-C*** illustrate the associations between the average deviation scores (ADSs) of brain morphological features and psychological levels of the dual-systems model. The ADS for surface thickness (ST) (β = 0.0406, SE = 0.009, *p* < 0.001) in the dorsolateral prefrontal cortex (DLPFC) was significantly associated with cognitive control, whereas the ADS for DLPFC surface area (SA) was not (β = -0.000604, SE = 0.0091, *p* = 0.947). Meanwhile, the ADS for subcortical volume (SV) of ventral striatum was significantly associated with sensation-seeking, as shown in *Panel C*.

***Panels D–E*** illustrate the trajectory of the imbalance score, calculated as the difference between the ventral striatum (VS) subcortical volume ADS and the DLPFC ST ADS, with scores ranging from -4 to 4. Higher values represent greater maturity of the socioemotional system compared to the cognitive-control system. The GAMLSS model fit for the imbalance trajectory, shown in Panel D, demonstrates negative scores during adolescence, gradually decreasing over time, reflecting the development of cognitive control. Panel E presents decreasing variance across the sample during adolescence, and Panel F indicates a decline in the growth rate of the imbalance.

### FIGURE S4. The Global Imbalance Trajectory and Model Fit.

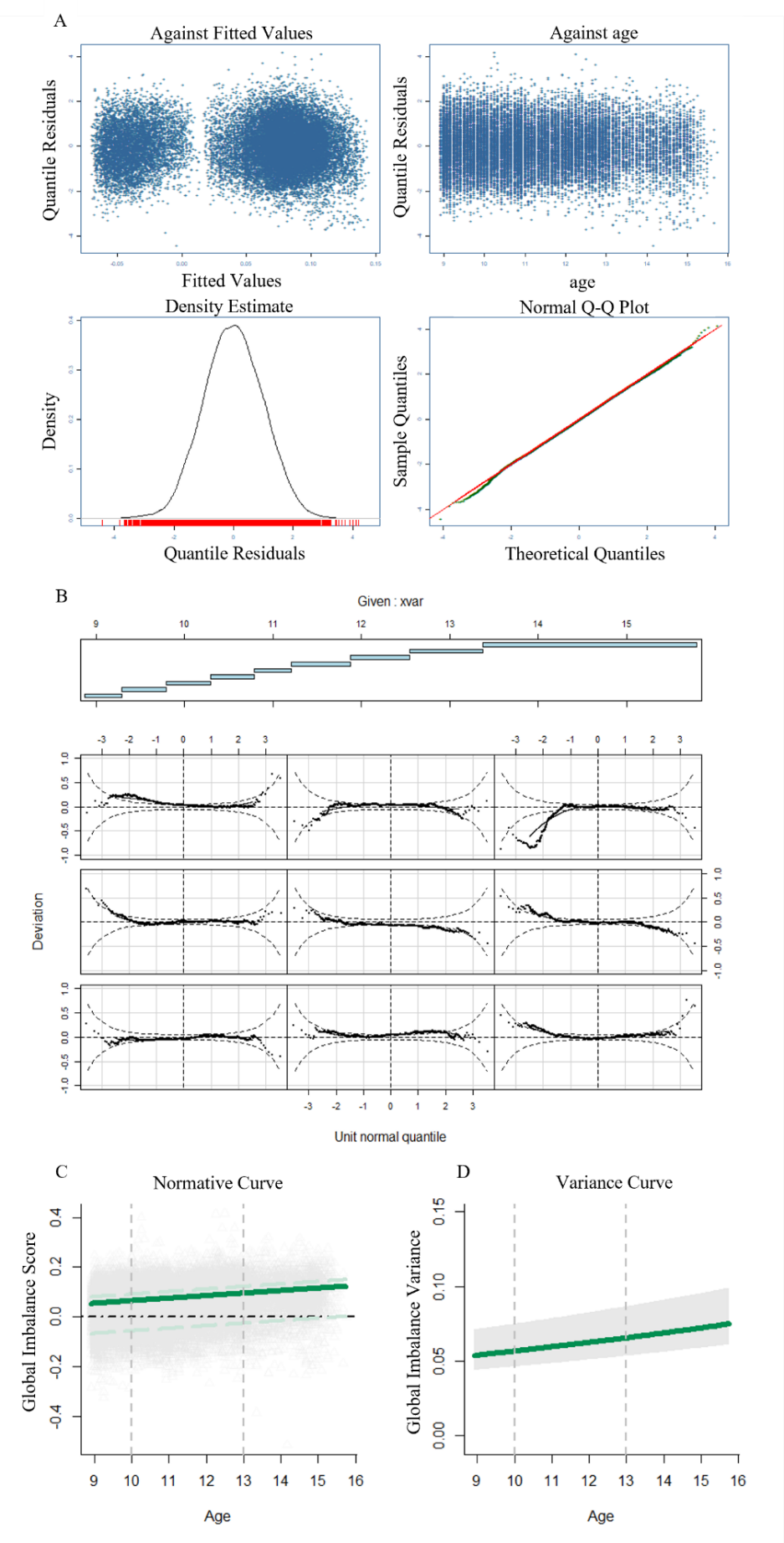

### FIGURE S5. The Alternative Imbalance Trajectory and Model Fit.

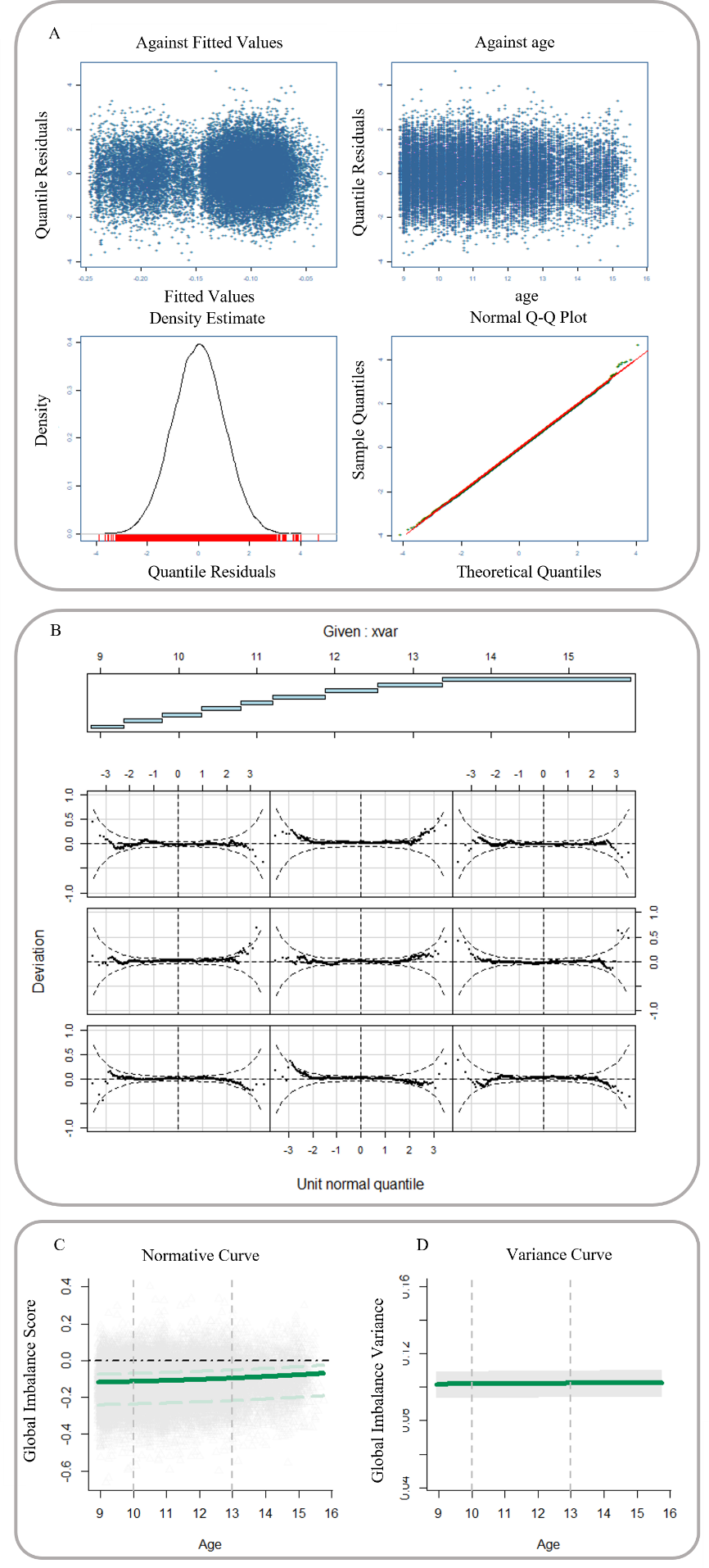

### TABLE S2. The Relationships Between the Broad Externalizing and the Imbalance Scores in Segmented Polynomial Models.

| **Segmented Polynomial models** | **Imbalance score, 1** | **Imbalance score, 2** | **Imbalance score, 3** | **Imbalance score, 4** | **Imbalance score, 5** |
| --- | --- | --- | --- | --- | --- |
| 2 | **B = 21.25, SE = 8.714, *p* = 0.014*** | **B = -13.74, SE = 8.274, *p* = 0.097.** | - | - | - |
| 3 | **B = 21.30, SE = 8.714, *p* = 0.014*** | **B = -13.75, SE = 8.274, *p* = 0.097.** | B = -6.146, SE = 8.200, *p* = 0.453 | - | - |
| 4 | **B = 21.30, SE = 8.714, *p* = 0.014*** | **B = -13.75, SE = 8.275, *p* = 0.097.** | B = -6.146, SE = 8.200, *p* = 0.453 | B = -0.392, SE = 8.197, *p* = 0.962 | - |
| 5 | **B = 21.27, SE = 8.713, *p* = 0.014*** | **B = -13.73, SE = 8.274, *p* = 0.097.** | B = -6.174, SE = 8.199, *p* = 0.451 | B = 0.434, SE = 8.197, *p* = 0.957 | **B = 13.71, SE = 8.201, *p* = 0.095.** |

### TABLE S3. The Model Selections in the Relationships with the Broad Externalizing Problems in GAMs.

| **GAMs** | **Term: Imbalance score** | **Term: Age** | **Term: Sex (male)** | **Term: Imbalance score * sex** | **Term: Site** | **Model Comparison** |
| --- | --- | --- | --- | --- | --- | --- |
| Reduced model | **B = 1.800, SE = 0.768,**  ***p* = 0.019*** | **B = -0.526, SE = 0.137,**  ***p* < 0.001***** | **B = 0.716, SE = 0.173,**  ***p* < 0.001***** | - | *edf* = 0.003, *F* = 0.003,  *p* = 0.316 | - |
| Model 2 | ***edf* = 2.851, *F* = 2.909,**  ***p* = 0.025*** | **B = -0.527, SE = 0.137,**  ***p* < 0.001***** | **B = 0.724, SE = 0.173,**  ***p* < 0.001***** | - | *edf* = 0.042, *F* = 0.044,  *p* = 0.309 | *F* (2.696, 9363) = 2.575,  *p =* 0.058 (.vs reduced model) |
| Model 3 | **B = 2.190, SE = 0.799,**  ***p* = 0.006**** | **B = -0.491, SE = 0.138,**  ***p* < 0.001***** | **B = 0.711, SE = 0.172,**  ***p* < 0.001***** | - | ***F* (21, 9339.7) = 3.367,**  ***p* < 0.001***** | ***F* (20.994, 9363) = 3.418, *p <* 0.001***** (.vs reduced model) |
| Model 4 | **B = 0.049, SE = 0.017,**  ***p* = 0.004**** | **B = -0.011, SE = 0.003,**  ***p* < 0.001***** | **B = 0.015, SE = 0.004,**  ***p* < 0.001***** | - | ***F* (21, 9339.7) = 3.367,**  ***p* < 0.001***** | **GCV = 0.0311** (.vs model 3) |
| Model 5 | **B = 0.049, SE = 0.017,**  ***p* = 0.004**** | ***edf =* 1.003*, F* = 13.2,**  ***p* < 0.001***** | **B = 0.015, SE = 0.004,**  ***p* < 0.001***** | - | ***F* (21, 9339.7) = 3.367,**  ***p* < 0.001***** | ***F* (0.006, 9342) = 0.289,**  ***p =* 0.018*** (.vs model 4) |
| Model 6 | *-* | ***edf =* 1.002*, F* = 12.847,**  ***p* < 0.001***** | **B = 0.015, SE = 0.004,**  ***p* < 0.001***** | Male*: edf =*1.001*, F* = 1.061,  *p* = 0.303  Female: *edf =* 2.106*, F* = 1.061,  *p* = 0.003** | ***F* (21, 9339.7) = 3.367,**  ***p* < 0.001***** | *F* (2.723, 9342) = 2.680,  *p =* 0.051 |

### TABLE S4. The Relationships Between the Broad Internalizing and the Imbalance Scores in Segmented Polynomial Models.

| **Segmented Polynomial models** | **Imbalance score, 1** | **Imbalance score, 2** | **Imbalance score, 3** | **Imbalance score, 4** | **Imbalance score, 5** |
| --- | --- | --- | --- | --- | --- |
| 2 | B = -14.79, SE = 9.060, *p* = 0.103 | **B = 23.79, SE = 8.574, *p* = 0.005**** | - | - | - |
| 3 | B = -14.88, SE = 9.060, *p* = 0.101 | **B = 23.80, SE = 8.574, *p* = 0.005**** | B = 8.281, SE = 8.493, *p* = 0.329 | - | - |
| 4 | B = -14.88, SE = 9.060, *p* = 0.100 | **B = 23.75, SE = 8.575, *p* = 0.005**** | B = 8.282, SE = 8.493, *p* = 0.329 | B = 7.428, SE = 8.490, *p* = 0.381 | - |
| 5 | B = -14.88, SE = 9.061, *p* = 0.100 | **B = 23.75, SE = 8.575, *p* = 0.005**** | B = 8.281, SE = 8.493, *p* = 0.329 | B = 7.430, SE = 8.491, *p* = 0.381 | B = 0.548, SE = 8.497, *p* = 0.948 |

### TABLE S5. The Model Selections in the Relationships With the Broad Internalizing Problems in GAMs.

| **GAMs** | **Term: Imbalance score** | **Term: Age** | **Term: Sex (male)** | **Term: Imbalance score * sex** | **Term: Site** | **Model Comparison** |
| --- | --- | --- | --- | --- | --- | --- |
| Reduced model | B = -1.447, SE = 0.802,  *p* = 0.071. | B = 0.364, SE = 0.143,  *p* < 0.011* | B = 0.829, SE = 0.180,  *p* < 0.001*** | - | *edf* = 0.729, *F* = 2.692,  *p* = 0.054. | - |
| Model 2 | *edf* = 2.778, *F* = 3.701,  *p* = 0.008** | B = 0.365, SE = 0.143,  *p* = 0.011* | B = 0.814, SE = 0.180,  *p* < 0.001*** | - | *edf* = 0.740, *F* = 2.830,  *p* = 0.051. | *F* (2.538, 9359) = 4.839,  *p =* 0.004** |
| Model 3 | *edf* = 2.757, *F* = 3.312,  *p* = 0.014* | B = 0.449, SE = 0.144,  *p* = 0.002** | B = 0.839, SE = 0.179,  *p* < 0.001*** | - |  | *F* (20.043,9339.5) = 4.719, *p <* 0.001*** |
| Model 4 | *edf =* 2.708*, F* = 3.451,  *p* = 0.011∗ | B = 0.008, SE = 0.003,  *p* = 0.003** | B = 0.018, SE = 0.004,  *p* < 0.001*** | - |  | - |
| Model 5 | *edf =* 2.711*, F* = 3.450,  *p* = 0.012∗ | *edf =* 1.002*, F* = 8.601,  *p* = 0.003** | B = 0.018, SE = 0.004,  *p* < 0.001*** | - |  | *F* (0.007,9340) = 0.767,  *p =* 0.019* |
| Model 6 | *edf =* 2.744*, F* = 3.486,  *p* = 0.011* | *edf =* 1.000*, F* = 8.690,  *p* = 0.003** | - | Male*: edf =* 0.015*, F* = 0.225,  *p* = 0.936  Female: *edf =* 1.001*, F* = 1.688,  *p* = 0.194 |  | *F* (1.073,9338.5) = 1.937,  *p =* 0.163 |

### TABLE S6. The Relationships Between the Broad Externalizing and the Imbalance Scores in Segmented Polynomial Models.

| **Segmented Polynomial models** | **Imbalance score, 1** | **Imbalance score, 2** | **Imbalance score, 3** | **Imbalance score, 4** | **Imbalance score, 5** |
| --- | --- | --- | --- | --- | --- |
| 2 | **B = 35.08, SE = 8.981, *p* < 0.001***** | B = -12.01, SE = 8.313, *p* = 0.148 | - | - | - |
| 3 | **B = 34.91, SE = 8.980, *p* < 0.001***** | B = -12.03, SE = 8.312, *p* = 0.148 | B = 11.53, SE = 8.206, *p* = 0.160 | - | - |
| 4 | **B = 34.94 SE = 8.980, *p* < 0.001***** | B = -12.08, SE = 8.313, *p* = 0.146 | B = 11.46, SE = 8.207, *p* = 0.162 | B = 6.678, SE = 8.182, *p* = 0.414 | - |
| 5 | **B = 34.94, SE = 8.981, *p* < 0.001***** | B = -12.08, SE = 8.313, *p* = 0.146 | B = 11.46, SE = 8.207, *p* = 0.162 | B = 6.682, SE = 8.183, *p* = 0.414 | B = -39.83, SE = 8.184, *p* = 0.961 |

### TABLE S7. The Relationships Between the Broad Externalizing and the Imbalance Scores in GAMs.

| **GAMs** | **Term: Imbalance score** | **Term: Age** | **Term: Sex (male)** | **Term: Imbalance score * sex** | **Term: Site** | **Model Comparison** |
| --- | --- | --- | --- | --- | --- | --- |
| Reduced model | **B = 2.934, SE = 0.758,**  ***p* < 0.001***** | **B = -0.562, SE = 0.137,**  ***p* < 0.001***** | **B = 0.740, SE = 0.172,**  ***p* < 0.001***** | - | *edf* = 0.003, *F* = 0.003,  *p* = 0.315 | - |
| Model 2 | **B = 3.345, SE = 0.830,**  ***p* < 0.001***** | **B = -0.522, SE = 0.138,**  ***p* < 0.001***** | **B = 0.737, SE = 0.172,**  ***p* < 0.001***** | - |  | ***F* (20.994, 9363) = 3.383,**  ***p <* 0.001*** (.vs reduced model)** |
| Model 3 | ***edf* = 1.761, *F* = 7.857,**  ***p* < 0.001***** | **B = -0.521, SE = 0.138,**  ***p* < 0.001***** | **B = 0.742, SE = 0.172,**  ***p* < 0.001***** | - |  | *F* (1.241, 9342) = 1.798, *p =* 0.178 (.vs model 2) |
| Model 4 | **B = 0.070, SE = 0.018,**  ***p* < 0.001***** | **B = -0.011, SE = 0.003,**  ***p* = 0.003**** | **B = 0.015, SE = 0.003,**  ***p* < 0.001***** | - |  | - |
| Model 5 | **B = 0.070, SE = 0.018,**  ***p* < 0.001***** | ***edf =* 1.002*, F* = 14.91,**  ***p* < 0.001***** | **B = 0.015, SE = 0.003,**  ***p* < 0.001***** | - |  | ***F* (0.005,9342) = 0.263,**  ***p =* 0.018*** |
| Model 6 | - | ***edf =* 1.003*, F* = 13.996,**  ***p* < 0.001***** | - | Male*: edf =* 1.448*, F* = 3.115,  *p* = 0.034*  Female: *edf =* 1.004*, F* = 5.666,  *p* = 0.017* |  | - |

### TABLE S8. The Relationships Between the Broad Internalizing and the Alternative Imbalance Scores in Segmented Polynomial Models.

| **Segmented Polynomial models** | **Imbalance score, 1** | **Imbalance score, 2** | **Imbalance score, 3** | **Imbalance score, 4** | **Imbalance score, 5** |
| --- | --- | --- | --- | --- | --- |
| 2 | B = -14.79, SE = 9.060, *p* = 0.102 | B = 23.79, SE = 8.574, *p* = 0.005** | - | - | - |
| 3 | B = -14.88, SE = 9.060, *p* = 0.101 | B = 23.80, SE = 8.574, *p* = 0.005** | B = 8.281, SE = 8.493, *p* = 0.329 | - | - |
| 4 | - | - | - | - | - |
| 5 | - | - | - | - | - |

### TABLE S9. The Relationships Between the Broad Externalizing and the Alternative Imbalance Scores in GAMs.

| **GAMs** | **Term: Imbalance score** | **Term: Age** | **Term: Sex (male)** | **Term: Imbalance score * sex** | **Term: Site** | **Model Comparison** |
| --- | --- | --- | --- | --- | --- | --- |
| Reduced model | B = -1.272, SE = 0.791,  *p* = 0.107 | B = 0.383, SE = 0.143,  *p* = 0.007** | B = 0.843, SE = 0.179,  *p* < 0.001*** | - | *edf* = 0.740, *F* = 2.856,  *p* = 0.049* | - |
| Model 2 | *edf* = 2.337, *F* = 2.639,  *p* = 0.0459* | B = 0.380, SE = 0.143,  *p* = 0.008** | B = 0.834, SE = 0.179,  *p* < 0.001*** | - | *edf* = 0.743, *F* = 2.897,  *p* = 0.048* | *F* (1.990, 9362.1) = 3.739,  *p =* 0.024* (.vs reduced model) |
| Model 3 | *edf* =2.342, *F* = 2.198,  *p* = 0.084 | B = 0.455, SE = 0.144,  *p* = 0.002** | B = 0.852, SE = 0.179,  *p* < 0.001*** | - |  | *F* (20.075, 9360) = 4.711, *p <* 0.001*** (.vs model 2) |
